# Dynamic regulation of the COPII interactome and collagen trafficking by site-specific glycosylation of Sec24D

**DOI:** 10.1101/2025.06.13.659590

**Authors:** Tetsuya Hirata, Dharmendra Choudhary, Brittany J. Bisnett, Erik J. Soderblom, Ela W. Knapik, Michael Boyce

**Author notes:** Correspondence: Tetsuya Hirata, Ph.D., and Michael Boyce, Ph.D., Department of Biochemistry, Duke University School of Medicine, Durham, NC 27710, USA. Lead contact: Michael Boyce, Ph.D.

## Abstract

Coat protein complex II (COPII) mediates anterograde trafficking from the endoplasmic reticulum (ER). While the core COPII machinery is well-characterized, how cells regulate COPII to accommodate large cargoes, including collagens, remains incompletely understood. Here, we show that the cargo-selecting COPII subunit Sec24D is modified by site-specific O-linked β-*N*-acetylglucosamine (O-GlcNAc) in its N-terminal intrinsically disordered region upon induction of collagen transport. These glycosylations are required for collagen trafficking in human cells and developing zebrafish. Crosslinking proteomics demonstrated that each O-GlcNAcylation influences the Sec24D interactome in a distinct way, revealing novel mediators of COPII function. In particular, Sec24D glycosylation is required for its interaction with myoferlin, which unexpectedly facilitates fusion of ER exit sites (ERES) and the ER-Golgi intermediate compartment (ERGIC) to enable collagen transport. Our results establish Sec24D O-GlcNAcylation as a dynamic regulator of COPII protein-protein interactions and collagen trafficking and identify myoferlin as a novel mediator of this process.

## Introduction

The secretory pathway is required for the trafficking and function of approximately one-third of the eukaryotic proteome^1,2^. In humans, dysfunction of intracellular protein transport causes severe diseases, including neurological, skeletal, and hematological disorders^3,4^. Coat protein complex II (COPII) is an essential and highly conserved multiprotein complex mediating anterograde protein trafficking from the endoplasmic reticulum (ER) to the ER-Golgi intermediate compartment (ERGIC) or Golgi. COPII carriers assemble at ER exit cites (ERESs) from five core components, Sar1, Sec23, Sec24, Sec13, and Sec31, with Sec23-Sec24 heterodimers forming an inner coat and Sec13-Sec31 heterotetramers an outer coat^1–4^. In animals, COPII trafficking also relies on accessory proteins, such as Sec16A, Sec23IP, and TANGO1 and its homologs^5,6^. Decades of rigorous studies have illuminated the formation and structures of COPII carriers using genetic, biochemical, and cell biological approaches^1,7–10^. However, significant aspects of COPII trafficking remain incompletely understood, such as how transport is dynamically regulated to accommodate physiological stimuli, stresses, and unusually large cargoes. For example, mature collagen trimers exiting the ER are ∼300-500 nm rods, whereas the original model of COPII carriers was a spherical vesicle of ∼60-80 nm diameter. Genetic evidence clearly shows that collagen transport is COPII-dependent^11–19^, raising the question of how the coat and its accessory proteins adapt to manage the trafficking challenge of large cargoes.

Post-translational modifications (PTMs) are likely a general mode of COPII regulation. Indeed, COPII proteins are decorated with a variety of PTMs, including phosphorylation, ubiquitination, and glycosylation^20,21^. We have studied COPII protein modification by O-GlcNAc, or O-linked β-*N*-acetylglucosamine, an abundant and essential PTM in humans and myriad other eukaryotes^22–24^. O-GlcNAcylation is the sole form of intracellular glycosylation in mammals, with ∼5,000 proteins and ∼7,000 glycosylation sites reported in humans to date^25^. O-GlcNAc cycling is governed by the essential enzymes O-GlcNAc transferase (OGT), which adds GlcNAc moieties to Ser/Thr residues of substrates, and O-GlcNAcase (OGA), which removes them^22–24^. This rapid reversibility affords the dynamic regulation of protein function in response to a wide range of stimuli^22–24,26–29^. We previously used mass spectrometry (MS) to identify O-GlcNAc sites on multiple COPII components, including Sec23A, Sec31A, Sec24C, and Sec24D^30–32^, and we and others have reported functional roles for the O-GlcNAcylation of Sec23A, Sec31A, and Sec24C in COPII formation and protein transport^30–33^. However, the impact and the regulation of O-GlcNAcylation on other COPII proteins remains unexplored.

In the present work, we focused on Sec24D, one of four cargo-selecting Sec24 paralogs in vertebrates^1,3,4,34^. Mutations in the human *SEC24D* gene cause subtypes of Cole-Carpenter Syndrome (CCS) and osteogenesis imperfecta (OI)^11–14^, both skeletal disorders associated with the abnormal formation, reduced mass, and increased fragility of bones resulting from decreased production or secretion of collagen^35^. Similar collagen trafficking defects and skeletal dysmorphology are observed in the *bulldog* zebrafish and *vertebra imperfecta* medaka mutants, which harbor nonsense mutations in *Sec24d*, and in Sec24D loss-of-function morphant zebrafish^15,16^. Therefore, vertebrate Sec24D is the essential Sec24 paralog for collagen transport *in vivo*. Previously, we identified six O-GlcNAcylation sites of human Sec24D by MS^30^ (Fig. 1A). Interestingly, three glycosites are in the N-terminal intrinsically disordered region (IDR), the domain least conserved among the Sec24 paralogs. The IDRs of other COPII components are proposed to regulate their protein-protein interactions and activities, particularly in the outer coat^36–38^, but the function of the Sec24D IDR remains enigmatic. Given its key role in cargo selection and collagen trafficking in health and disease^1,39–42^, understanding the regulation of Sec24D is an important goal.

**Figure 1.**
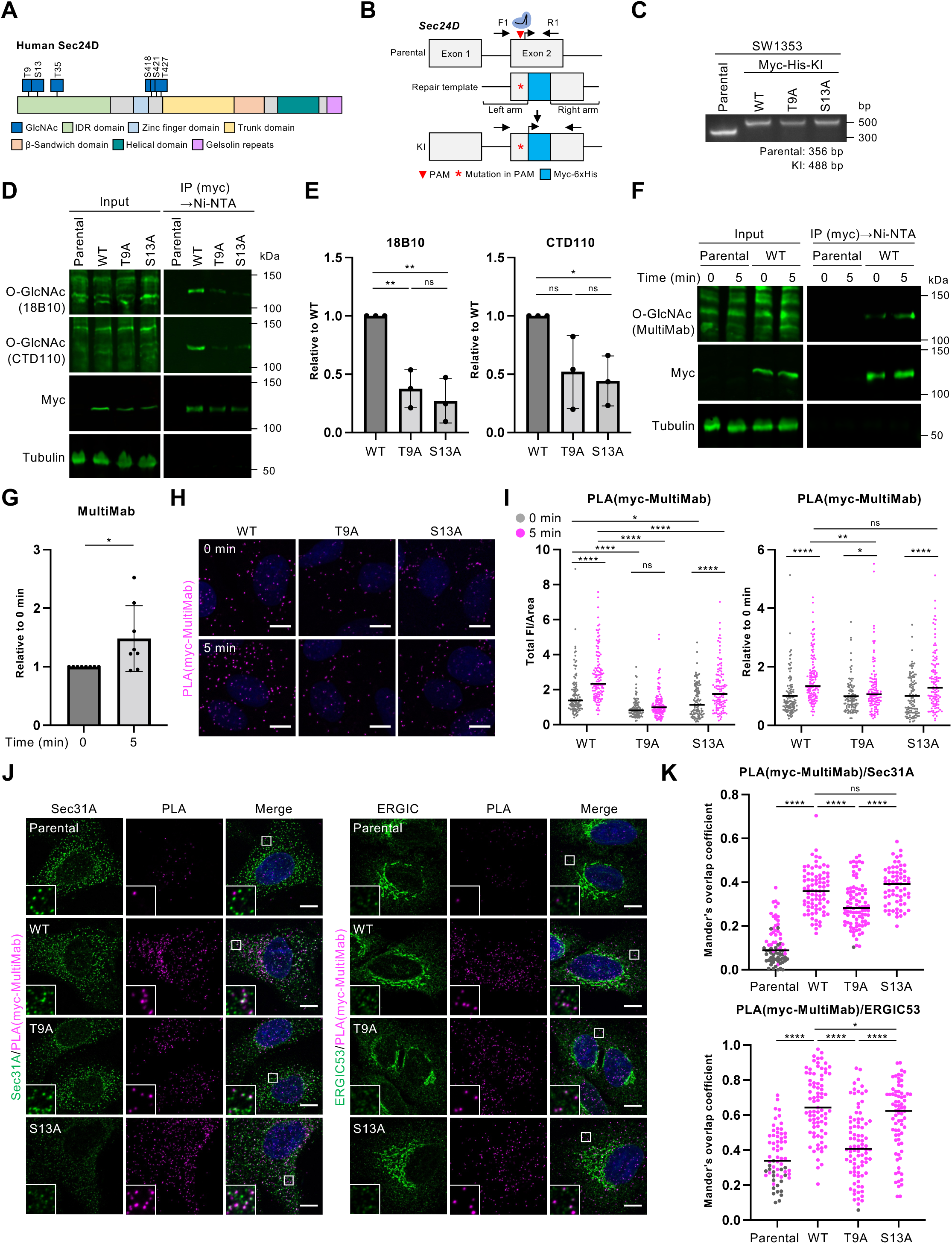
Sec24D O-GlcNAcylation on T9 is induced upon collagen transport. A. Domain structure and O-GlcNAc sites of human Sec24D. B. CRISPR strategy to generate *Sec24D* knock-in cells. C. PCR genotyping of Sec24D^WT^, Sec24D^T9A^, and Sec24D^S13A^ SW1353 single cell-derived clones, using primers depicted in B. D. Myc-6xHis tagged endogenous Sec24D was purified from Sec24D^WT^, Sec24D^T9A^, and Sec24D^S13A^ cells by tandem myc IP and denaturing Ni-NTA affinity and analyzed by IB. E. Quantification of experiments depicted in D. Signal from O-GlcNAc blots of purified Sec24D was normalized to myc signal, and intensities relative to Sec24D^WT^ were calculated (n = 3). Data were analyzed by one-way ANOVA with *post hoc* Tukey test. * *p* < 0.05; ** *p* < 0.01; ns: not significant. F. Myc-6xHis tagged endogenous Sec24D was purified from Sec24D^WT^, Sec24D^T9A^, and Sec24D^S13A^ cells by tandem myc IP and denaturing Ni-NTA affinity at 0 or 5 minutes after collagen release and analyzed by IB. G. Quantification of experiments depicted in F. Signal from O-GlcNAc blots of purified Sec24D was normalized to myc signal, and intensities relative to time-zero were calculated (n = 8). Data were analyzed by Welch’s *t*-test. * *p* < 0.05. H. O-GlcNAcylated Sec24D was visualized by PLA in Sec24D^WT^, Sec24D^T9A^, and Sec24D^S13A^ cells at 0 or 5 minutes after collagen release. Scale bars: 10 µm. I. Quantification of experiments depicted in H. Left: Total fluorescence intensity (FI) of PLA signal normalized to per-cell area (n = 3). Right: Total FI, normalized to cell area at time-zero (n = 3). Data were analyzed by two-way ANOVA with *post hoc* Tukey test. * *p* < 0.05; ** *p* < 0.01; **** *p* < 0.0001; ns: not significant. J. Parental SW1353, Sec24D^WT^, Sec24D^T9A^, and Sec24D^S13A^ cells were analyzed by PLA for O-GlcNAcylated Sec24D (myc-MultiMab antibodies), and Sec31A or ERGIC53 was detected by IF. Scale bars: 10 µm. K. Quantification of experiments depicted in J. Mander’s overlap coefficients (MOCs) between PLA signals and Sec31A or ERGIC53 was calculated (N = 3). MOC data with Costes *p* value < 0.95 were deemed to be statistically indistinguishable from random overlap and are displayed as gray dots. One-way ANOVA with *post-hoc* Tukey test was conducted as statistical analysis. * *p* < 0.05; **** *p* < 0.0001; ns: not significant.

Here, we report the functions of O-GlcNAc on T9 and S13 in the Sec24D IDR. Using biochemical and cell biological approaches, we show that dynamic O-GlcNAcylation at these sites regulates COPII-dependent collagen trafficking in cultured human cells and developing zebrafish embryos. Crosslinking proteomics demonstrated that the T9 and S13 glycosites influence distinct aspects of the Sec24D interactome, identifying both known and novel COPII components, cargoes, and accessory proteins whose associations with Sec24D rely on IDR glycosylation. Finally, we show that one such novel O-GlcNAc-dependent Sec24D interactor, myoferlin (MYOF), mediates the fusion of ERES and ERGIC membranes to facilitate collagen transport. Our work provides new insight into the dynamic regulation of COPII function in response to collagen cargo demand and identifies MYOF as novel functional mediator of early secretory pathway trafficking.

## Results

### O-GlcNAcylation on T9 is dynamically regulated upon collagen transport

Among the six identified O-GlcNAc sites on Sec24D (Fig. 1A), we focused on T9 and S13 for further investigation because they are well conserved among vertebrates (Fig. S1), and they reside in the poorly understood IDR. We used CRISPR methods^43^ to edit the *SEC24D* locus in human SW1353 chondrosarcoma cells, introducing an N-terminal myc-6xHis tag and either wild type (WT) sequence or a T9A or S13A mutation, thereby eliminating each glycosite individually (Fig. 1B). SW1353 cells secrete endogenous collagen and are an established human cell system to study large cargo trafficking ^31,44,45^. Successful homozygous editing in single cell-derived clones was verified via PCR (Fig. 1B-C) and Sanger sequencing (Fig. S2A-B). Expression and localization of myc-tagged Sec24D^WT^, Sec24D^T9A^, and Sec24D^S13A^ in the respective clones were confirmed by immunoblot (IB) and immunofluorescence imaging (IF) (Fig. S2C-D). Importantly, the protein expression of Sec24A, B, and C was comparable in all three cell lines (Fig. S2E-F).

Quantitative anti-O-GlcNAc IBs with two different monoclonals showed that glycosylation is significantly reduced on the Sec24D^T9A^ and Sec24D^S13A^ mutants, relative to Sec24D^WT^, identifying T9 and S13 as major O-GlcNAc sites in live cells (Fig. 1D-E). (Multiple anti-O-GlcNAc monoclonal antibodies are commonly used in IB and IF experiments because these reagents are known to exhibit amino acid sequence preferences^46^.) We next asked whether T9 or S13 glycosylation responds to collagen trafficking as a physiological stimulus. Following a well-established procedure^47^, Sec24D^WT^, Sec24D^T9A^, or Sec24D^S13A^ cells were incubated at a non-permissive temperature for three hours, to cause collagen retention in the ER, and then shifted to a permissive temperature and supplemented with ascorbate, a cofactor for proline hydroxylation and collagen maturation^48^, to trigger a synchronous release of collagen from the ER via COPII. O-GlcNAcylation of Sec24D^WT^ was significantly increased 5 minutes after collagen release (Fig. 1F-G), suggesting that Sec24D is dynamically glycosylated in response to collagen transport.

COPII trafficking initiates at ERES in response to cargo demand^1–4^. To determine whether collagen transport triggers site-specific Sec24D O-GlcNAcylation at ERES and/or other functionally relevant locations, we established a proximity ligation assay (PLA)^49^ for glycosylation of endogenous, CRISPR-tagged Sec24D using anti-myc and anti-O-GlcNAc antibodies (Fig. S3A-B). Our PLA was dependent on myc-tagged Sec24D expression, and signal was increased by treatment with Thiamet-G (an OGA inhibitor) and decreased by Ac_4_5SGlcNAc (5SG; an OGT inhibitor), demonstrating the specificity of the assay (Fig. S3A-B). Next, we measured Sec24D O-GlcNAc levels in Sec24D^WT^, Sec24D^T9A^, and Sec24D^S13A^ cells after collagen release. Consistent with our IB data (Fig. 1F-G), PLA signal on Sec24D^WT^ increased 5 minutes after collagen release (Fig. 1H-I). Interestingly, the glycosite mutants had subtly different defects in collagen-induced glycosylation: Both mutants exhibited diminished O-GlcNAcylation, relative to Sec24D^WT^, at both time-points, but Sec24D^S13A^ still induced O-GlcNAcylation from 0 to 5 min, whereas Sec24D^T9A^ O-GlcNAcylation was insensitive to cargo release (Fig. 1H-I). These results indicate that T9 O-GlcNAcylation is dynamically induced by collagen transport, whereas S13 glycosylation is likely constitutive.

We next examined the subcellular location of O-GlcNAcylated Sec24D. Glycosylated Sec24D^WT^ PLA signal colocalized significantly with Sec31A and ERGIC53, markers of ERES/COPII carriers and ERGIC, respectively (Fig. 1J-K), indicating that O-GlcNAcylated Sec24D is largely incorporated into COPII carriers and localized at ERES and the ERGIC. In Sec24D^T9A^ cells, PLA colocalization with Sec31A and ERGIC53 was significantly decreased (Fig. 1J-K), perhaps due to inefficient recruitment of Sec31A to ERESs and/or defective protein transport (see below). Taken together, these results demonstrate that collagen trafficking induces site-specific IDR O-GlcNAcylation on pools of ERES- and ERGIC-localized Sec24D.

### O-GlcNAcylation of T9 and S13 has distinct roles in collagen transport

Our results suggested that O-GlcNAcylation of Sec24D at T9 and S13 might be functionally important for collagen transport. To test this hypothesis, we first examined intracellular collagen accumulation. Sec24D^WT^, Sec24D^T9A^, and Sec24D^S13A^ cells were treated with ascorbate for 5 hours, and levels of intracellular collagen were quantified. When transport is efficient, little to no collagen is retained intracellularly under these conditions^19,31,50,51^. Indeed, intracellular collagen I α1 (COL1A1) was barely detectable in Sec24D^WT^ cells (Fig. 2A-B). By contrast, COL1A1 significantly accumulated in the ER of Sec24D^T9A^ and Sec24D^S13A^ cells, indicating a transport defect (Fig. 2A-B). To determine whether Sec24D IDR glycosylation influences collagen transport kinetics, cells were subjected to temperature-shift to induce synchronous collagen release and analyzed at different time points. As expected, in Sec24D^WT^ cells, COL1A1 was within the ER at 0 minutes and moved towards the Golgi in a time-dependent manner (Fig. 2C-D). By contrast, both Sec24D^T9A^ and Sec24D^S13A^ cells showed COL1A1 retention in the ER even 15 minutes after cargo release, demonstrating that collagen transport was significantly delayed (Fig. 2C-D). Though expression levels of Sec24D^T9A^ or Sec24D^S13A^ were somewhat lower than that of Sec24D^WT^ (Fig. S2F), the observed trafficking defects were not caused by Sec24D protein levels *per se* because supraphysiological expression of Sec24D^WT^ rescued the intracellular collagen accumulation of Sec24D^T9A^ or Sec24D^S13A^ cells, whereas overexpression of Sec24D^T9A^ or Sec24D^S13A^ did not (Fig. S4A-B). These results demonstrate that O-GlcNAc on T9 and S13 is essential for efficient collagen transport from the ER to the Golgi.

**Figure 2.**
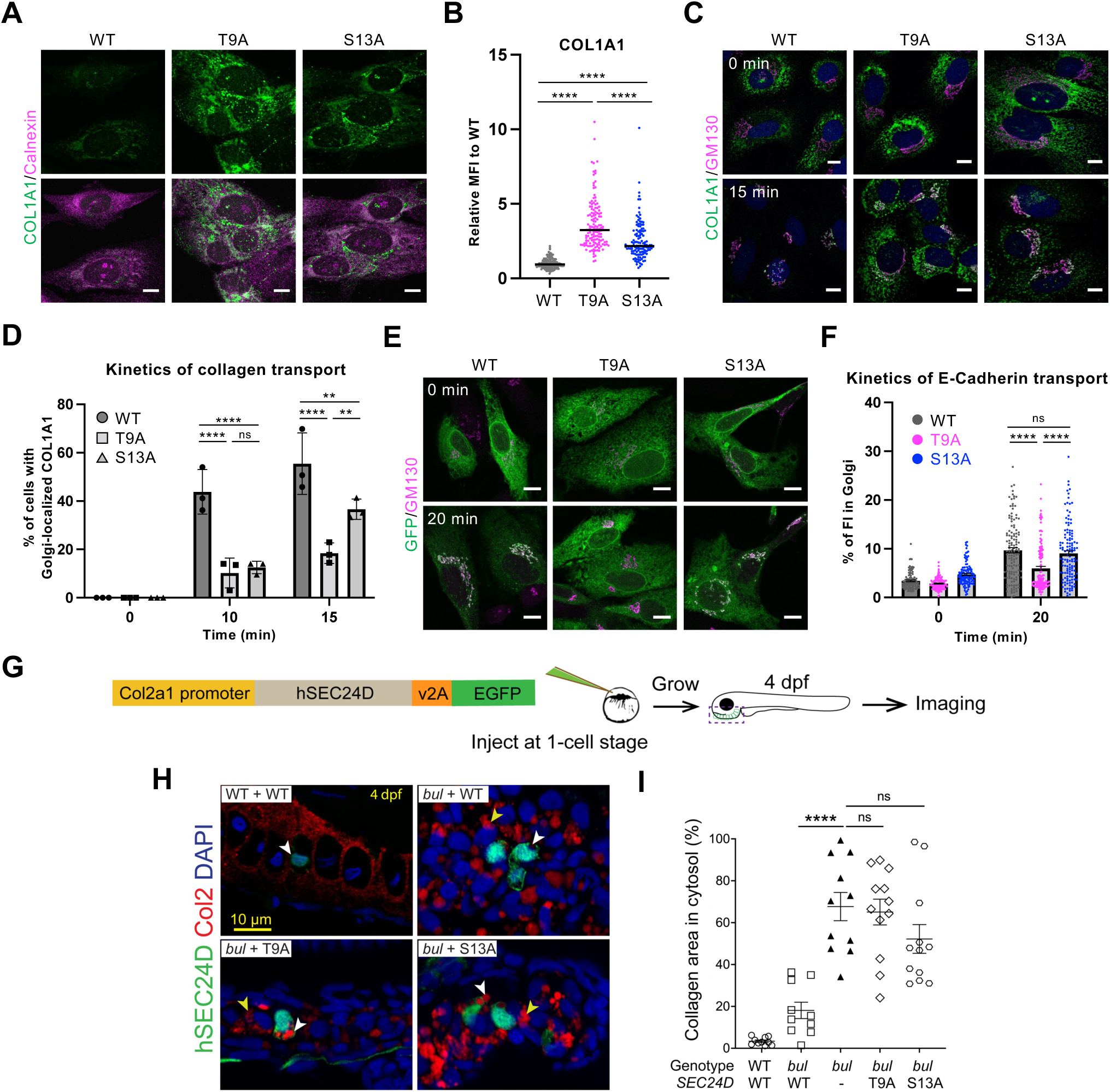
O-GlcNAcylation of T9 and S13 is required for collagen transport in cultured human cells and developing zebrafish. A. Sec24D^WT^, Sec24D^T9A^, and Sec24D^S13A^ cells were treated with 50 µg/ml sodium ascorbate for 5 h and analyzed by IF for collagen I (COL1A1) and calnexin, an ER marker. Scale bars: 10 µm. B. Quantification of experiments depicted in A. Mean fluorescence intensity (MFI) of COL1A1, normalized to Sec24D^WT^, was calculated (n = 3). Data were analyzed by one-way ANOVA with *post hoc* Tukey test. **** *p* < 0.0001. C. Sec24D^WT^, Sec24D^T9A^, and Sec24D^S13A^ cells were incubated at 40 °C for 3 h, shifted to 32 °C plus 50 µg/ml sodium ascorbate and 100 µg/ml cycloheximide for the indicated times, and analyzed by IF for COL1A1 and GM130 (Golgi marker). Scale bars: 10 µm. D. Quantification of experiments depicted in C. Percentage of cells with Golgi-localized COL1A1 was calculated (n = 3). Data were analyzed by two-way ANOVA with *post hoc* Tukey test. ** *p* < 0.01; **** *p* < 0.0001; ns: not significant. E. Sec24D^WT^, Sec24D^T9A^, and Sec24D^S13A^ cells expressing the E-cadherin RUSH system were incubated with 50 µM biotin and 100 µg/ml cycloheximide for the indicated times and analyzed by IF for E-cadherin (GFP) and GM130. Scale bars: 10 µm. F. Quantification of experiments depicted in E. Percentage of GFP FI co-localizing with the Golgi (GM130) was calculated (n = 3). Data were analyzed by two-way ANOVA with *post hoc* Sidak’s test. **** *p* < 0.0001; ns: not significant. G. Experimental design for mosaic expression of human *SEC24D* (hSEC24D) gene in zebrafish rescue experiments. H. WT or *bulldog* (*bul*) zebrafish embryos were injected with expression constructs encoding human SEC24D^WT^, SEC24D^T9A^, or SEC24D^S13A^ and were analyzed by IF at 4 dpf. White arrowheads indicate hSEC24D-GFP-expressing cells (green). WT fish show no intracellular collagen retention. hSEC24D^WT^ expression largely rescues the collagen retention in *bul* fish, whereas hSEC24D^T9A^- or hSEC24D^S13A^-injected *bul* fish retain intracellular collagen (red). Yellow arrowheads indicate retained intracellular collagen in neighboring non-hSEC24D-expressing cells. I. Quantification of experiments depicted in H. Intracellular collagen area was calculated (n = WT: 10; *bul*+hSEC24D^WT^: 10; *bul:* 11; *bul*+hSEC24D^T9A^: 12; *bul*+hSEC24D^S13A^: 12 cells) from a minimum of 6 different embryos in each group, and data were analyzed by one-way ANOVA with Dunnett’s multiple comparison test. **** *p* < 0.0001; ns: not significant. Error bars are standard error of the mean.

Collagen is an unusually large cargo and utilizes unconventional COPII carriers^52–55^. To test whether Sec24D O-GlcNAcylation on T9 and S13 is also required for the transport of smaller COPII clients, we took advantage of the model cargo E-cadherin^56,57^ and the retention using selective hook (RUSH) system^58^. In a RUSH assay, the cargo is tagged with a streptavidin-binding peptide (SBP) and is retained in the ER via co-expression of a streptavidin-KDEL fusion as a hook^58^. Biotin added to the culture medium out-competes SBP binding to streptavidin, leading to the synchronous release of the cargo from the ER^58^. In Sec24D^WT^ cells, the E-cadherin RUSH probe was retained in the ER at time zero, and its Golgi localization increased by 20 minutes after biotin addition (Fig. 2E-F). In contrast, E-cadherin trafficking was significantly delayed in Sec24D^T9A^ cells but not in Sec24D^S13A^ cells at the 20-min time-point (Fig. 2E-F). These results suggest that O-GlcNAcylation on T9 and S13 may have distinct roles in protein trafficking, with S13 O-GlcNAcylation being required for collagen transport, whereas T9 O-GlcNAcylation may be needed for a wider range of cargoes.

### O-GlcNAcylation of Sec24D is required for collagen transport in vivo

We next examined whether Sec24D T9 and S13 O-GlcNAcylation is required for collagen trafficking *in vivo*. We injected WT or *Sec24d*-deficient *bulldog* mutant zebrafish with Tol2 constructs encoding human SEC24D^WT^, SEC24D^T9A^, and SEC24D^S13A^, driven by the zebrafish collagen 2α1 promoter (Fig. 2G). Importantly, O-GlcNAc signaling and the T9 and S13 glycosites are conserved in zebrafish Sec24D (Fig. S1)^31,59–61^. Next, we quantified intracellular collagen accumulation in *bulldog* and WT chondrocytes^15^. WT chondrocytes had less than 5% cell area occupied by collagen, compared to 68% in *bulldog* chondrocytes (Fig. 2H-I). Expression of SEC24D^WT^ fully rescued this retention defect, reducing intracellular collagen occupancy to 18% in *bulldog* fish (Fig. 2H-I). However, neither SEC24D^T9A^ nor SEC24D^S13A^ produced substantial rescue, with both glycosite mutants being statistically indistinguishable from the Sec24D-deficient *bulldog* mutant (Fig. 2H-I). These results indicate that Sec24D O-GlcNAcylation on T9 and S13 has a conserved and essential function in collagen transport *in vivo*.

### Sec24D O-GlcNAcylation regulates its interaction with COPII components and novel mediators of COPII function

We next sought to understand the molecular mechanisms underlying the defective collagen transport in Sec24D^T9A^ and Sec24D^S13A^ cells. The IDRs of some COPII components mediate their interactions with binding partners^36–38^, and O-GlcNAcylation is well-known to influence protein-protein interactions as well^31,62,63^. Therefore, we hypothesized that Sec24D IDR glycosylation may influence its interactome, and we tested this notion in Sec24D^WT^, Sec24D^T9A^, and Sec24D^S13A^ cells. Protein transport is rapid, and most protein-protein interactions between Sec24D and cargoes or other interactors are likely to be transient or weak. To capture such interactions, we treated Sec24D^WT^, Sec24D^T9A^, and Sec24D^S13A^ cells with the cell-permeable chemical crosslinker dithiobis[succinimidyl propionate] (DSP), stringently purified Sec24D and crosslinked proteins through tandem myc immunoprecipitation (IP) and denaturing Ni-NTA affinity, and reductively cleaved the DSP crosslinks to liberate interacting proteins. We visualized proteins co-purifying with Sec24D by SDS-PAGE and silver stain. Multiple bands were detected across all tagged cell lines, whereas untagged parental cells showed very low background, demonstrating the utility of this approach (Fig. S5A).

We performed shotgun MS proteomics with label-free quantitation over three biological replicates to profile the Sec24D interactomes in Sec24D^WT^, Sec24D^T9A^, and Sec24D^S13A^ cells. We identified 360 proteins as specific interactors of Sec24D^WT^ (i.e., enriched relative to untagged, negative-control parental cell samples), including strong enrichment of proteins annotated as involved in ER to Golgi vesicle-mediated transport, the COPII system, actin cytoskeleton organization, Golgi organization, and retrograde vesicle-mediated transport (Fig. 3A-B). In particular, known COPII coat components and accessory proteins were confirmed as Sec24D^WT^ interactors (Fig. 3A and Fig. S5B and Table S1). Many other proteins related to transport, such as COPI coat proteins, cargo receptors, SNAREs, and small GTPases, were also enriched in Sec24D^WT^ samples (Fig. 3A and S5B and Table S1). These results demonstrate that functionally relevant Sec24D interactors can be identified by our approach. In addition, we identified novel candidate Sec24D^WT^-interacting proteins with no previously reported roles in ER-Golgi transport, such as components of the PI3 kinase and Hippo signaling pathways (Fig. S5B and Table S1). These data provide a new catalog of the Sec24D interactome from a human cell type secreting endogenous collagen, an obligate Sec24D cargo of pathophysiological interest.

**Figure 3.**
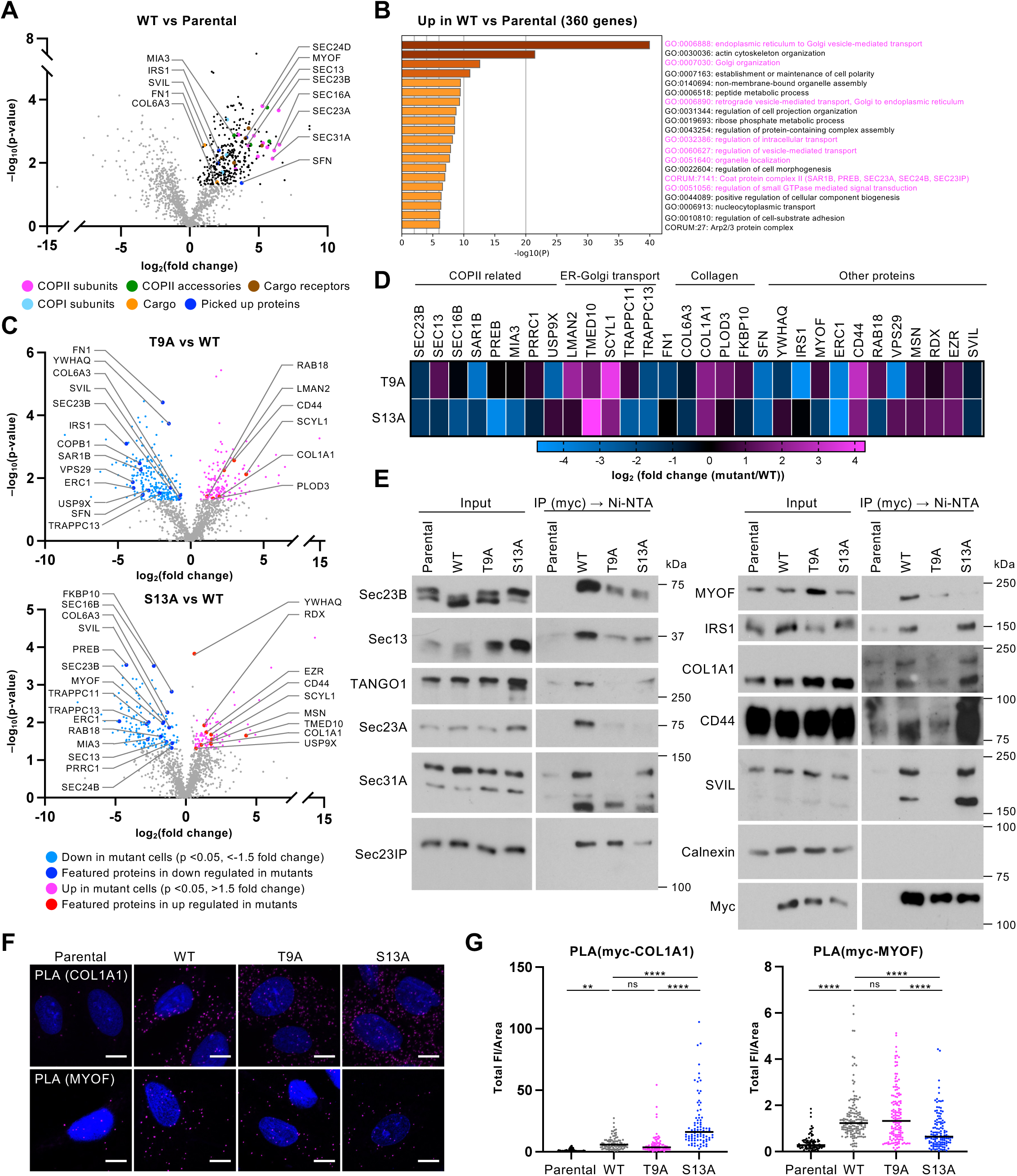
Sec24D O-GlcNAcylation regulates its interaction with COPII components and novel trafficking accessory proteins. A. Parental SW1353 or Sec24D^WT^ cells were treated with DSP, and myc-6xHis tagged endogenous Sec24D^WT^ was tandem purified by myc IP and denaturing Ni-NTA purification. After reductive crosslinker cleavage, interacting proteins were analyzed by liquid-chromatography (LC)-MS/MS analyses. Proteins with ≥ 1.5 fold change and *p* value < 0.05 in Sec24D^WT^ versus parental samples from three independent biological samples were considered significantly enriched. Volcano plot shows 360 specific interactors of Sec24D^WT^. Proteins not meeting the above criteria are displayed as gray dots. B. Gene ontology analysis of significantly enriched proteins identified in experiments depicted in A. C. Interacting proteins of myc-6xHis-tagged endogenous Sec24D^T9A^ and Sec24D^S13A^ were purified and analyzed as in A, relative to Sec24D^WT^ interactors. Volcano plots show 147 and 246 interactors significantly enriched and depleted, respectively, in Sec24D^T9A^ samples, and 92 and 168 proteins enriched or depleted, respectively, in Sec24D^S13A^ samples, relative to Sec24D^WT^. Proteins not meeting the above criteria are displayed as gray dots. D. Heatmap depicting selected proteins that demonstrated altered interactions with Sec24D^T9A^ or Sec24D^S13A^, relative to Sec24D^WT^, in MS data. E. Myc-6xHis-tagged endogenous Sec24D was purified from Sec24D^WT^, Sec24D^T9A^, and Sec24D^S13A^ cells without DSP crosslinking by tandem myc IP and non-denaturing Ni-NTA affinity and analyzed by IB. F. Parental SW1353, Sec24D^WT^, Sec24D^T9A^, and Sec24D^S13A^ cells were analyzed via IF and PLA for Sec24D-COL1A1 interaction (myc-COL1A1 antibodies) and Sec24D-MYOF interaction (myc-MYOF antibodies). Scale bars: 10 µm. G. Quantification of experiments depicted in F. Total FI of PLA was normalized to cell area (n = 3). Data were analyzed by one-way ANOVA with *post hoc* Tukey test. ** *p* < 0.01; **** *p* < 0.0001; ns: not significant.

Next, we examined whether the Sec24D^WT^ interactome differed from those of Sec24D^T9A^ and Sec24D^S13A^. Relative to Sec24D^WT^, 147 and 246 proteins exhibited significantly increased and decreased interactions, respectively, with Sec24D^T9A^, and 92 and 168 proteins exhibited significantly increased and decreased interactions with Sec24D^S13A^ (Fig. S5C). Notably, we did not detect COL1A1 as a Sec24D^WT^ interactor, but it was identified in both Sec24D^T9A^ and Sec24D^S13A^ samples (Fig. 3C-D and Table S1), likely because the transport defect in mutant cells results in a longer dwell time for Sec24D-collagen interactions. Procollagen-lysine,2-oxoglutarate 5-dioxygenase 3 (PLOD3), which catalyzes both proline hydroxylation and glycosylation on collagen^64,65^, and FKBP10, a chaperone specific for pro-collagen^66–68^, were also enriched in the Sec24D^T9A^ interactome, relative to Sec24D^WT^. Intriguingly, many COPII components and subunits of the TRAPP complex (e.g., TRAPPC11 and TRAPPC13), which tethers ER-derived transport carriers to the ERGIC or Golgi, were significantly depleted from the Sec24D^T9A^ and Sec24D^S13A^ interactomes, relative to Sec24D^WT^ (Fig. 3C-D and Fig. S5D). These results suggest that COPII formation, cargo capture and release, and/or carrier fusion with target membranes may be defective in Sec24D mutant cells.

To confirm our proteomics results, we first performed tandem non-denaturing IP/Ni-NTA purification of Sec24D^WT^ from uncrosslinked samples and analyzed candidate interactors by IB. These results corroborated the MS data, demonstrating associations between Sec24D^WT^ and COPII components and novel interactors, including myoferlin (MYOF), supervillin (SVIL), insulin receptor substrate 1 (IRS1), and CD44, and 14-3-3 proteins (Fig. 3E and S5E). Interactions between Sec24D^T9A^ or Sec24D^S13A^ and COPII-related proteins, such as Sec23A, Sec23B, Sec31A, Sec13, Sec23IP, and TANGO1 (encoded by *MIA3*) were generally reduced, relative to Sec24D^WT^ (Fig. 3E). Associations with novel interactors were also altered in Sec24D^T9A^ and Sec24D^S13A^ cells. For example, the interaction with MYOF was decreased for Sec24D^S13A^, whereas associations with COL1A1, IRS1, CD44, and SVIL were decreased for Sec24D^T9A^ but increased for Sec24D^S13A^ (Fig. 3E). To confirm and extend these results, we established highly specific PLAs for Sec24D and selected interactors (Fig. 3F-G). The *in cellulo* Sec24D-COL1A1 and Sec24D-MYOF interactions were markedly increased and decreased, respectively, for Sec24D^S13A^, relative to Sec24D^WT^ (Fig. 3F-G), consistent with our MS and IP/IB data. Collectively, these results indicate that O-GlcNAcylation on both T9 and S13 is necessary for functional Sec24D protein-protein interactions but that the two sites may have distinct mechanistic roles in regulating these associations and downstream cargo transport.

### Sec24D O-GlcNAcylation is required for recruitment of COPII components to ERESs

The glycosite mutants of Sec24D exhibited reduced interaction with COPII components (Fig. 3C-E and S5D), suggesting at least two possible causes: 1) Sec24D^T9A^ and Sec24D^S13A^ might not properly localize to ERESs, or 2) Recruitment of other COPII components to ERES might be inefficient in Sec24D^T9A^ and Sec24D^S13A^ cells. In testing the first possibility, IF revealed that co-localization of Sec24D^T9A^ or Sec24D^S13A^ and the ERES marker Sec16A was higher than for Sec24D^WT^ (Fig. 4A-B), indicating that glycosite mutant recruitment to ERES is not reduced but that productive COPII carrier formation at ERES may be inhibited. Indeed, in testing the second possibility, IF demonstrated that co-localization of Sec24D^T9A^ and the outer coat protein Sec31A was significantly lower than for Sec24D^WT^ (Fig. 4C-D). Co-localization of Sec24D^S13A^ and Sec31A was slightly higher than that of Sec24D^WT^ and Sec31A (Fig. 4C-D). However, in Sec24D^S13A^ cells, the mean fluorescence intensity (MFI) of Sec31A puncta was significantly decreased, compared to Sec24D^WT^ cells (Fig. 4C-D), even though total protein levels of Sec31A were comparable across all cell lines (Fig. S6A-B), indicating abnormal Sec31A recruitment to ERES in Sec24D^S13A^ cells. These results suggest that O-GlcNAcylation on T9 and S13 of Sec24D is necessary for the recruitment of the key COPII component Sec31A to ERESs or other functional sites, perhaps by independent mechanisms.

**Figure 4.**
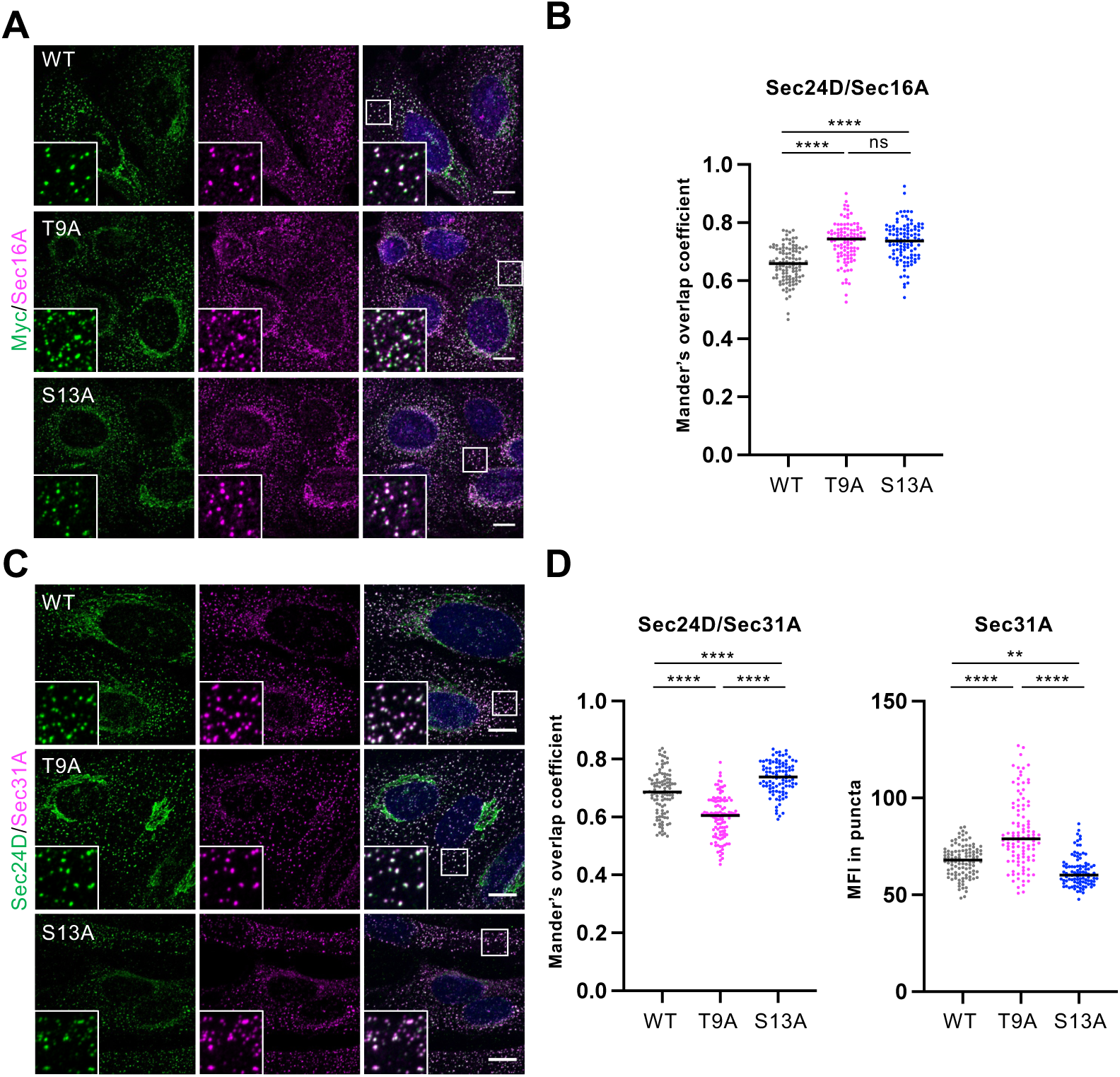
Sec24D O-GlcNAcylation is required for recruitment of COPII components to ERESs. A. Sec24D^WT^, Sec24D^T9A^, and Sec24D^S13A^ cells were analyzed by IF. Scale bars: 10 µm. B. Quantification of experiments depicted in A. Mander’s overlapping coefficient (MOC) between Sec24D (myc) and Sec16A (ERES marker) was calculated (n = 3). Data were analyzed by one-way ANOVA with *post hoc* Tukey test. **** *p* < 0.0001; ns: not significant. C. Sec24D^WT^, Sec24D^T9A^, and Sec24D^S13A^ cells were analyzed by IF. Scale bars: 10 µm. D. Quantification of experiments depicted in C. Left: MOC between Sec24D (myc) and Sec31A. Right: MFI of Sec31A puncta was quantified (n = 3). Data were analyzed by one-way ANOVA with *post hoc* Tukey test. ** *p* < 0.01; **** *p* < 0.0001; ns: not significant.

### MYOF is a novel mediator of collagen transport from the ER

Our MS proteomics and IB validation experiments identified several new interactors of Sec24D that might play previously unknown roles in collagen transport. We selected MYOF for further investigation as it is one of the top five proteins in the Sec24D^WT^ interactome, with respect to number of identified peptides (Fig. S5F and Table S1), and its interaction is dependent on Sec24D IDR O-GlcNAcylation (Fig. 3E-G). Interestingly, a high-throughput study of the human protein interactome also identified a potential interaction between MYOF and Sec24C, a closest paralog of Sec24D^69^, though this result has not been independently validated. MYOF is a type II transmembrane protein with a large cytosolic region containing seven C2 domains (C2A-C2G)^70,71^ (Fig. 5A). Its best-known function is membrane repair and fusion in myoblasts^72,73^, and it is reported to participate in the endocytic transport of the epidermal growth factor receptor^74,75^. However, prior to this study, MYOF had no known role in the early secretory pathway.

**Figure 5.**
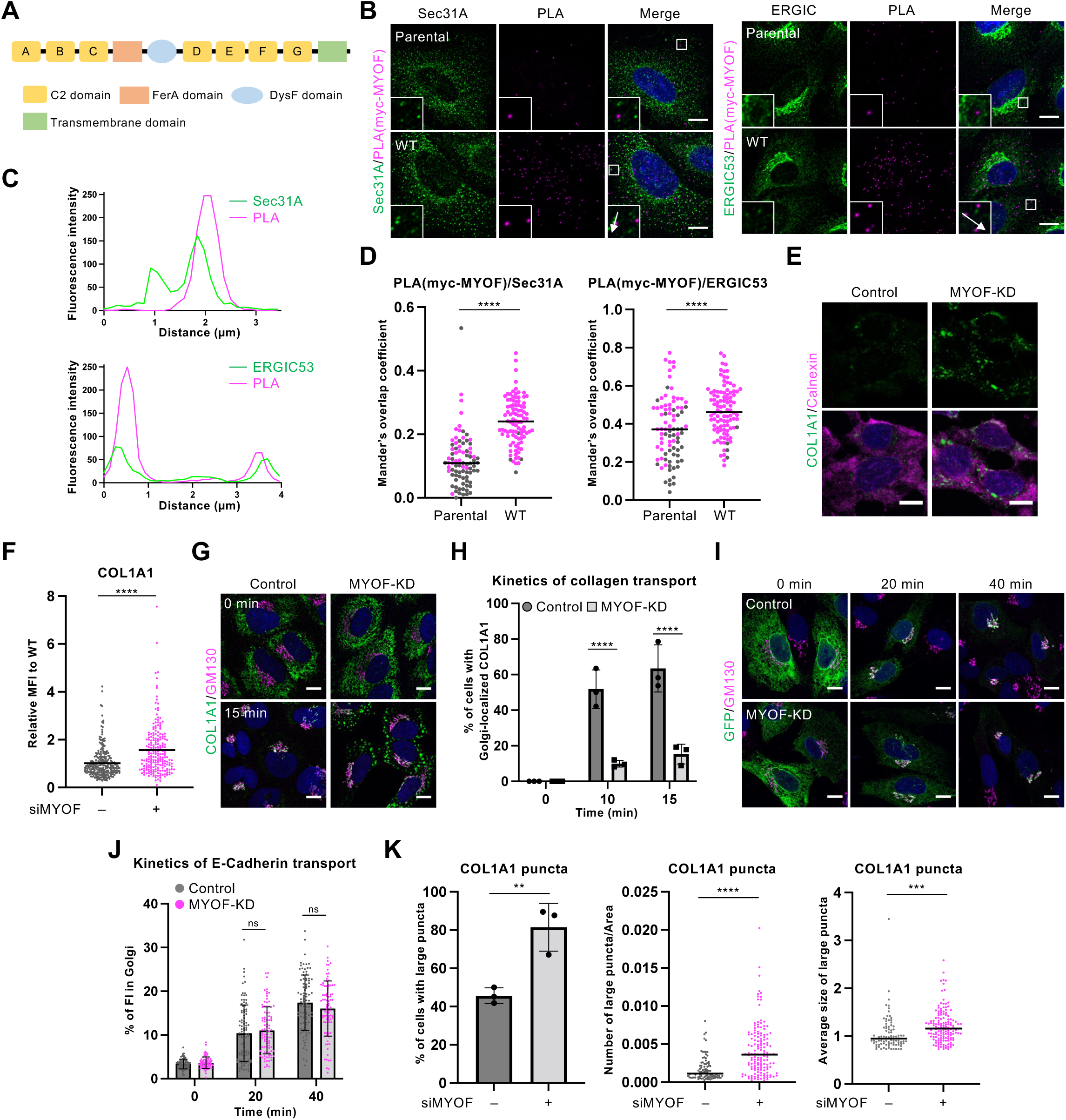
MYOF is a novel mediator of collagen transport. A. Domain structure of MYOF. B. Sec24D^WT^, Sec24D^T9A^, and Sec24D^S13A^ cells were analyzed by PLA for Sec24D-MYOF interaction (myc-MYOF antibodies) and Sec31A (left) or ERGIC53 (right) via IF. Scale bars: 10 µm. C. Line plots display the FIs of PLA and Sec31A (top) or ERGIC53 (bottom) signal along the inset arrow paths in B. D. From experiments depicted in B, MOCs between PLA and Sec31A (left) or PLA and ERGIC53 (right) were calculated (n = 3). Data were analyzed by unpaired Student’s or Welch’s *t*-test. MOC data with Costes *p* value < 0.95 were deemed to be statistically indistinguishable from random overlap and are displayed as gray dots. **** *p* < 0.0001. E. Control and MYOF-knockdown SW1353 cells were treated with 50 µg/ml sodium ascorbate for 5 h and then analyzed by IF. Scale bars: 10 µm. F. Quantification of experiments depicted in E. MFI of COL1A1 was measured and normalized to the control sample (n = 3). Data were analyzed by Welch’s *t*-test. **** *p* < 0.0001. G. Control and MYOF-knockdown SW1353 cells were incubated at 40 °C for 3 h, shifted to 32 °C plus 50 µg/ml sodium ascorbate and 100 µg/ml cycloheximide for the indicated times, and analyzed by IF. Scale bars: 10 µm. H. Quantification of experiments depicted in G. Percentage of cells with Golgi-localized COL1A1 in each sample was calculated (n = 3). Data were analyzed by two-way ANOVA with *post hoc* Sidak’s test. **** *p* < 0.0001. I. Control and MYOF-knockdown SW1353 cells expressing the E-cadherin RUSH system were incubated with 50 µM biotin and 100 µg/ml cycloheximide for the indicated times and analyzed by IF. Scale bars: 10 µm. J. Quantification of experiments depicted in I. Percentage of COL1A1 FI in the Golgi was calculated for each sample (n = 3). Data were analyzed by two-way ANOVA with *post hoc* Sidak’s test. ns: not significant. K. Quantification of experiments depicted in G at 15 min time-points. Left: Percentage of cells with large COL1A1 puncta (defined as ≥ 0.7 µm^2^ size) was calculated for each sample (n = 3). Data were analyzed by unpaired Student’s *t*-test. Middle: Average size of COL1A1 puncta was calculated (n = 3). Data were analyzed by unpaired Student’s *t*-test. Right: Number of large COL1A1 puncta was calculated and normalized to cell area (n = 3). Data were analyzed by Welch’s *t*-test. ** *p* < 0.01; *** *p* < 0.001; **** *p* < 0.0001.

MYOF is reported to localize to the plasma membrane and multiple intracellular organelles, including the Golgi, endosomes, and lysosomes^76,77^. There is no previous experimental evidence of MYOF at ERESs or the ERGIC, the likely sites of its interaction with Sec24D, but a prediction tool for protein location at intracellular organelles^78^ suggested that MYOF may localize to the ERGIC (Fig. S7A-C). Specifically, the proteomic enrichment profile of MYOF resembled the averaged profile of endosomal and ERGIC/cis-Golgi resident proteins but not those of ERES proteins, and the 20 proteins with enrichment profiles most similar to MYOF include endosomal and ERGIC/cis-Golgi proteins (Fig. S7C-H). Indeed, PLA signal between Sec24D^WT^ and MYOF overlapped or closely co-localized with both Sec31A and ERGIC53 (Fig. 5B-D). These results indicate that Sec24D and MYOF interact in a location near to both ERESs and the ERGIC.

Next, we investigated the functional relevance of MYOF to collagen transport. CRISPR deletion of MYOF in SW1353 cells resulted in collagen expression changes at the mRNA and protein levels (Fig. S8A-C), confounding analysis of transport experiments. Therefore, we transiently depleted MYOF expression in SW1353 cells via siRNA knockdown (Fig. S9A-B) and examined intracellular collagen accumulation as before. Control cells displayed very low intracellular COL1A1, whereas MYOF-depleted cells showed significantly higher levels (Fig. 5E-F). Time-resolved experiments demonstrated that COL1A1 transport was severely delayed by MYOF depletion (Fig. 5G-H). These results indicate that MYOF is required for efficient ER-to-Golgi collagen transport. Finally, to determine whether the transport function of MYOF is specific to collagen, we carried out E-cadherin RUSH assays. E-cadherin transport was not significantly affected by MYOF depletion, suggesting a potentially specific function of MYOF in collagen transport (Fig. 5I-J).

### MYOF mediates ERES-ERGIC fusion to accommodate large COPII cargoes

Bulky collagens are thought to exit the ER in large, unconventional COPII carriers^6,53,54,79^, and previous studies reported that TANGO1-mediated fusion of ERES and ERGIC membranes is required for this process^6,50,51^. In our own transport experiments, we noticed that COL1A1 puncta were larger and more numerous in MYOF-depleted cells, compared to controls (Fig. 5E, 5G and 5K). Given these observations and the fact that the C2 domains of MYOF reportedly bind membrane^80,81^, we hypothesized that MYOF might contribute ERES-ERGIC fusion via membrane interactions. To test this hypothesis, we first characterized the localization of accumulated collagen puncta in MYOF-depleted cells. Sec31A was near COL1A1 in both control and MYOF-depleted cells, indicating that collagen accumulates at ERESs or COPII carriers upon MYOF-depletion (Fig. 6A). Furthermore, in control cells, ERGIC53 was detected at the edge of, and partially overlapping with, COL1A1 puncta (Fig. 6B-C, upper panels). By contrast, in MYOF-depleted cells, ERGIC53 signal was almost exclusively at the edge of COL1A1 puncta (Fig. 6B-C, lower panels). These observations support the model that MYOF participates in the fusion of ERES and ERGIC membranes during collagen transport.

**Figure 6.**
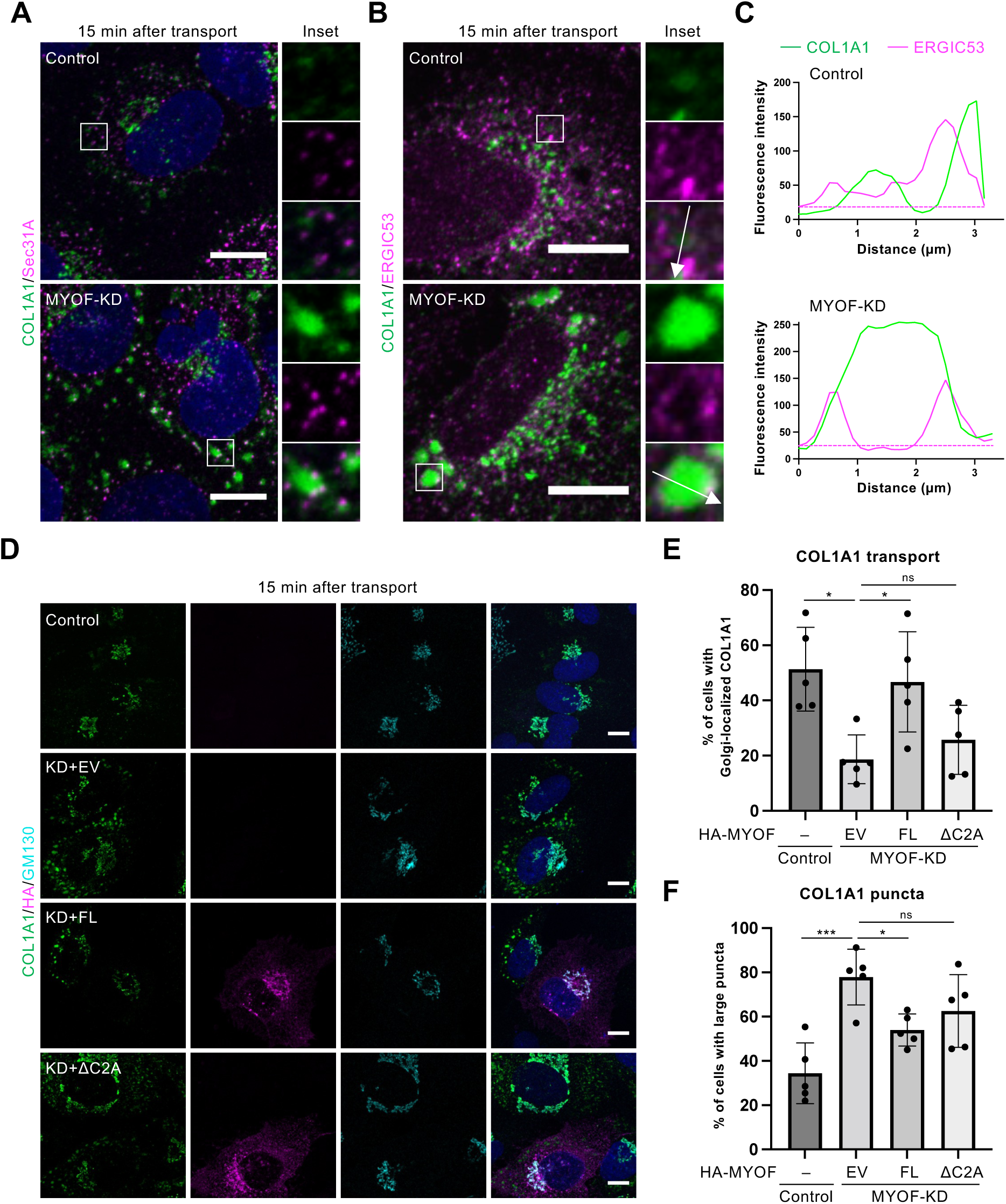
MYOF mediates ERES-ERGIC fusion to accommodate large COPII cargoes. A. Control and MYOF-knockdown SW1353 cells were incubated at 40 °C for 3 h, shifted to 32 °C plus 50 µg/ml sodium ascorbate and 100 µg/ml cycloheximide for 15 min, and analyzed by IF. Scale bars: 10 µm. B. Control and MYOF-knockdown SW1353 cells were incubated at 40 °C for 3 h, shifted to 32 °C plus 50 µg/ml sodium ascorbate and 100 µg/ml cycloheximide for 15 min, and analyzed by IF. Scale bars: 10 µm. C. Quantification of experiments depicted in B. Line plots display the FIs of COL1A1 and ERGIC53 signal along the inset arrow paths. Dotted lines represent the remaining noise in the ERGIC53 channel after background subtraction. D. MYOF-knockdown SW1353 cells were transfected with empty vector (EV) or siRNA-resistant expression constructs for full-length (FL) or ΔC2A mutant HA-MYOF. Cells were incubated at 40°C for 3 h, shifted to 32 °C plus 50 µg/ml sodium ascorbate and 100 µg/ml cycloheximide for 15 min, and analyzed by IF. Scale bars: 10 µm. E. Quantification of experiments depicted in D. Percentage of cells with Golgi-localized COL1A1 was calculated for each sample (n = 5). Data were analyzed by one-way ANOVA with *post hoc* Tukey test. * p < 0.05; ns: not significant. F. Quantification of experiments depicted in D. Percentage of cells with large COL1A1 puncta was calculated for each sample (n = 5). Data were analyzed by one-way ANOVA with *post hoc* Tukey test. * p < 0.05; *** p < 0.001; ns: not significant.

To further test this hypothesis, we conducted rescue experiments in MYOF-depleted cells by expressing siRNA-resistant full-length MYOF (FL) or a truncation mutant lacking the C2A domain (ΔC2A), which interacts with membranes in Ca^2+^-dependent manner^80^. Expression levels of FL and ΔC2A MYOF were equivalent, as judged by IB, and MYOF localization was not affected by C2A domain deletion (Fig. S10A-C). Re-expression of FL MYOF, but not ΔC2A, rescued the accumulation of large collagen puncta and transport delay of MYOF-depleted cells (Fig. 6D-F and Fig. S10D), demonstrating that the C2A domain of MYOF is required for collagen transport. Taken together, these results support the model that MYOF is required to facilitate the fusion of ERES and ERGIC membranes during collagen transport.

## Discussion

Collagens comprise up to 30% of the dry weight of the human body^82,83^, but how cells regulate and remodel the COPII machinery to accommodate such large cargoes remains incompletely understood. Our study indicates that dynamic, site-specific Sec24D IDR O-GlcNAcylation may play a key role in collagen transport in vertebrates. Based on our results, we propose the following stepwise model for Sec24D O-GlcNAcylation in collagen trafficking (Fig. 7): 1) S13 of Sec24D is constitutively O-GlcNAcylated and mediates recruitment of COPII components to ERESs. MYOF interacts directly or indirectly with S13-O-GlcNAcylated Sec24D. 2) T9 of Sec24D is rapidly O-GlcNAcylated in response to collagen maturation and transport demand. O-GlcNAc on T9 and S13 facilitates further COPII recruitment to ERESs to form carriers. MYOF binds ERES and ERGIC membranes. 3) MYOF facilitates ERES/ERGIC membrane fusion, permitting collagen egress from the ER.

**Figure 7.**
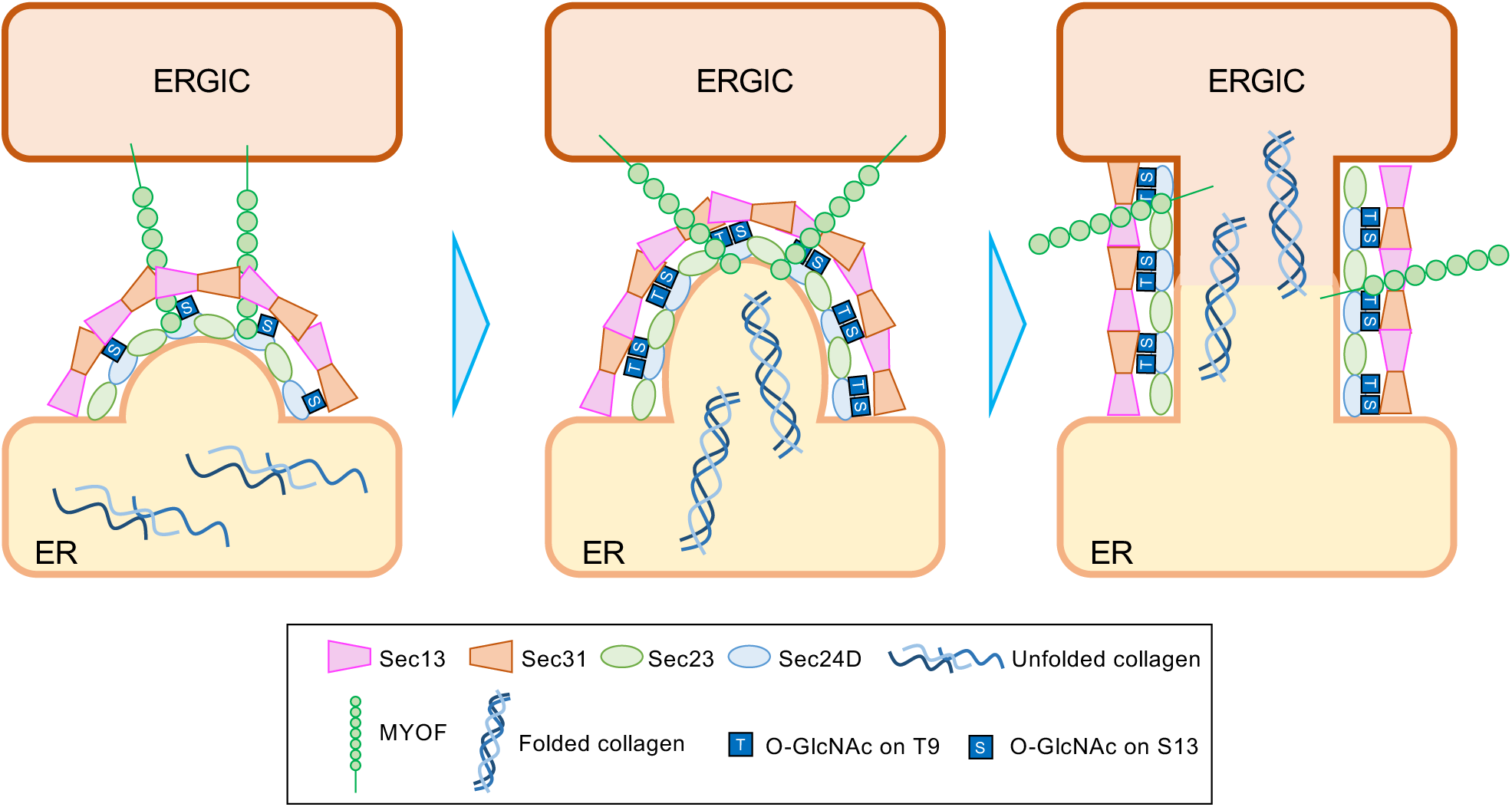
Working model of site-specific Sec24D IDR O-GlcNAcylation in collagen transport. Left: S13 of Sec24D is constitutively O-GlcNAcylated. MYOF interacts with O-GlcNAc-modified Sec24D. Middle: T9 of Sec24D is rapidly O-GlcNAcylated upon collagen transport. O-GlcNAc on T9 and S13 facilitates further recruitment of key interactors to ERESs, such as Sec23A, Sec23B, and Sec31A, generating large COPII carriers. MYOF binds the ER membrane at ERES via its C2A domain. Right: MYOF facilitates fusion of the ERGIC membrane with ERESs, mediating collagen egress from the ER. We note that variations on this model are also consistent with our data. For example, MYOF may reside in and/or bind the ERGIC or ER membrane or both, and Sec24D may interact with MYOF either directly or indirectly (i.e., in multiprotein complexes).

Our results demonstrate that the T9 and S13 Sec24D glycosites play important but distinct roles in collagen trafficking, with T9 O-GlcNAcylation likely required for a larger range of cargoes than S13 (Fig. 1F-K, 2A-F, and 3C-E). Further studies will be needed to dissect the mechanistic basis for the difference in T9-O-GlcNAc and S13-O-GlcNAc functions, but discrepancies in the interactomes of Sec24D^T9A^ and Sec24D^S13A^ may provide clues. For instance, MYOF interacts with Sec24D in an S13-O-GlcNAc-dependent manner (Fig. 3C-G), and its function may be specific to collagen transport (Fig. 5E-J). Conversely, the interaction of Sec24D with IRS1, a regulator of insulin signaling^84^, depends on T9-O-GlcNAc but not S13-O-GlcNAc (Fig. 3C-E). IRS1 has recently been reported to localize to the ER via interaction with vesicle-associated membrane protein-associated protein B and to undergo liquid-liquid phase separation (LLPSs) upon insulin stimulation^85^. Since COPII transport may also be regulated by LLPSs, as discussed below, IRS1 could participate in this process. Regardless of the most important downstream interactors or other mechanisms, it is notable that the first 20 amino acids of Sec24D are well-conserved across vertebrates, even though other regions of the IDR are more variable (Fig. S1), and neither Sec24D^T9A^ nor Sec24D^S13A^ rescued collagen trafficking in the *bulldog* zebrafish mutant (Fig. 2H-I). Therefore, our general model of Sec24D IDR O-GlcNAcylation in collagen trafficking (Fig. 7) may apply across a range of vertebrates.

The exact nature of COPII collagen carriers remains debated, with various reports describing large vesicles, tubules, or tunnels connecting the ER and ERGIC^9,10,86–88^. The assembly and stability of the inner coat is likely to be a main driver of COPII carrier morphology^34^, suggesting that regulation of Sec24D may play a role. In principle, our results are compatible with both vesicle and tubule/tunnel models, and Sec24D glycosylation may contribute to either process through a variety of potential mechanisms. For example, recent studies proposed that distinct forms of the COPII lattice mediate trafficking of general versus bulky cargo carriers^6,52^. Cryo-electron tomography of COPII on yeast microsomes suggested that COPII carrier morphology may be influenced by the inner coat, with tubules or spherical carriers alternately produced, depending on inner coat assembly and density^89^. Since Sec24D T9 and S13 O-GlcNAcylation is required for its interaction with other COPII subunits and for collagen transport (Fig. 2A-D and 3C-E), these PTMs might regulate COPII lattice formation and geometry by modulating the alignment of inner coat complexes. In addition, some COPII components, including Sec24D, undergo LLPS through multivalent protein-protein interactions of the IDR domains, and COPII transport is regulated by LLPSs^36^. Because O-GlcNAcylation is known to regulate LLPS formation in other contexts^90–92^, Sec24D IDR glycosylation may influence COPII carrier size and morphology through LLPS, an important hypothesis for future studies. Finally, Sec31A mono-ubiquitination by a CUL3-KLHL12 E3 ligase and other regulators, including ALG2, PEF1, and USP9X, is proposed to mediate the generation of large COPII carriers for bulky cargo transport and/or collagen degradation^53,54,93–95^. Interestingly, we identified the de-ubiquitination enzyme USP9X as an S13-influenced Sec24D interactor (Fig. 3C-D and Table S1). Since USP9X promotes the de-ubiquitination of PEF1 to inactivate the CUL3-KLHL12 E3 ligase, in turn decreasing the mono-ubiquitination of Sec31A^95^, the increased interaction of USP9X with Sec24D^S13A^ could indicate a potential mechanism to modulate COPII carrier size or fate.

We and others have previously shown that many COPII proteins are modified by O-GlcNAc^21,30–33^. Our current results are novel, in part, because they are the first report of rapid and dynamic O-GlcNAc change on a COPII subunit in response to cargo transport (Fig. 1F-I). An important question raised by our work is how the initiation of collagen transport triggers O-GlcNAc changes. One possibility is that transmembrane proteins communicate information about collagen assembly and recruitment at ERES to Sec24D and/or OGT. One such candidate mediator is TANGO1, which interacts with collagen and COPII via Sec23A^6^. Collagen binding by TANGO1 or its homologs might induce conformational changes in COPII components, such as Sec23A, Sec24D, or Sec31A, that recruit OGT to ERES, facilitating Sec24D O-GlcNAcylation. Sec23A itself is also O-GlcNAcylated^31^, and OGT was identified as a Sec24D interactor in our proteomics analysis (Table S1). Thus, either protein might recruit OGT in response to stimuli from the ER lumen. Another, mutually compatible possibility is that Sec24D O-GlcNAcylation is downstream of other PTM signaling cascades that respond to folded cargoes in the ER. For example, a recent report observed that the phosphorylation of over 70 proteins is changed within four minutes of collagen folding^57^, one minute earlier than our observation of increased Sec24D T9 O-GlcNAcylation (Fig. 1H-I). It is possible that collagen maturation induces a phosphorylation cascade that facilitates Sec24D O-GlcNAcylation by activating or recruiting OGT to ERESs or by inactivating or excluding OGA.

Our results also provide insight into the interactome of Sec24D and how it is influenced by site-specific IDR O-GlcNAcylation (Fig. 3). We anticipate that our Sec24D^WT^ interactome data will be a useful resource to the trafficking field, containing known and novel COPII accessory proteins, cargoes, regulators, and others (Fig. 3A). Our data also revealed – perhaps surprisingly – that the Sec24D interactome is significantly and differentially affected by mutation of either of two IDR glycosites (Fig. 3C-D). At present, it is still unclear how O-GlcNAcylation of the Sec24D IDR regulates protein-protein interactions. O-GlcNAc moieties can directly mediate protein-protein interactions^31,62^ and modulate protein folding^96^, suggesting that glycans on the IDR domain may recruit interactors through direct binding and/or by altering its secondary structure, important hypotheses to test in future work.

Our proteomics and other results also revealed MYOF to be a novel, glycosylation-dependent interactor of Sec24D with a previously unappreciated role in collagen trafficking (Fig. 3), though whether Sec24D and MYOF interact directly or indirectly (e.g., in a multiprotein complex) remains to be determined. While MYOF had no known function in the early secretory pathway, it was an attractive candidate for our follow-up experiments in part because it contains multiple C2 domains, which are well-known to bind membranes, and other proteins with multiple C2 domains, such as synaptotagmin-1, participate in anterograde protein transport^97^. The C2 and Fer A domains of MYOF are reported to bind phospholipids in a Ca^2+^-dependent manner *in vitro*^80,98^, and we demonstrated that the C2A domain is required for MYOF function in collagen transport (Fig. 6D-F). Moreover, we showed that MYOF interacts with Sec24D at the interface of ERESs and ERGIC membranes (Fig. 5B-D), supporting our proposal that MYOF recruits or tethers ERES and ERGIC membranes to facilitate their fusion (Fig. 7). Other ER-localized COPII components, tethering molecules, and SNARE proteins have been shown to mediate ERES/ERGIC membrane fusion, such as p125A (also known as Sec23IP), the NRZ complex (NAG, RINT1, and ZW10), syntaxin-18, and BNIP1^50,51,99,100^. MYOF might function together with these proteins or independently, via a parallel mechanism. Notably, MYOF may play a more specific role in the transport of large cargoes than other previously reported regulators because its depletion had no impact on E-cadherin transport (Fig. 5I-J), distinguishing it from other essential regulators of collagen trafficking, such as TANGO1, which is required for general COPII transport as well^55,101–105^. Future functional experiments on MYOF may further elucidate the regulatory and remodeling mechanisms that specifically accommodate COPII to bulky client proteins.

Finally, our results will motivate future investigations into the potential role of the Sec24D/O-GlcNAc/MYOF axis in human disease. The importance of *SEC24D* mutations in CCS and OI is well-established^11–14^, and future studies will examine the impact of these mutations and other skeletal disease states on Sec24D glycosylation. O-GlcNAcylation influences the differentiation of osteoblasts and osteoclasts, and dysregulated O-GlcNAc signaling is implicated in bone diseases such as osteoporosis and osteoarthritis^106–108^. Glycosylation of Sec24D may be involved in these contexts as well, though the crucial substrates and mechanisms are as-yet incompletely characterized. The potential role of MYOF in bone formation and maintenance also remains under-explored. No gross skeletal abnormalities have been reported in human patients with loss-of-function *MYOF* mutations or in *Myof*^-/-^ mice^72,73,109^. However, humans express four other Ferlin family members, in addition to MYOF, and dysferlin expression is upregulated in *Myof*^-/-^ mice^60^, suggesting that genetic compensation could obscure an *in vivo* role for MYOF in normal collagen trafficking in the skeleton. Indeed, our observation of COL1A1 expression changes in *MYOF*^-/-^ SW1353 cells is consistent with this hypothesis (Fig. S8). On the other hand, MYOF mutations do impact other tissues and cause human disease. Recently, a *MYOF* truncation mutation was reported to cause limb-girdle muscular dystrophy and cardiomyopathy in a human patient^73^, while a *MYOF* point mutation gave rise to arrhythmogenic right ventricular cardiomyopathy^109^, a rare autosomal dominant disorder causing severe arrhythmia and sudden cardiac death^110–112^. Interestingly, mutations in collagens *COL6A1*, *COL6A2*, *COL6A3*, and *COL12A1* are responsible for certain types of muscular dystrophy, including Ullrich syndrome and Bethlem myopathy^113^, and a *klhl40*-deficient zebrafish exhibits reduced SEC24D protein expression, defective collagen secretion, and muscle dysfunction^114^. Together, these data indicate specific connections between collagen secretion and muscle biogenesis or function, suggesting that collagen secretion defects may contribute to the pathogenesis of MYOF-deficient patients.

In summary, our studies have revealed new roles for both site-specific Sec24D IDR glycosylation and MYOF in collagen trafficking and have contributed to our knowledge of the COPII machinery through Sec24D interactome profiling. In these ways, our work addresses the dynamic regulation of COPII transport and the mechanisms of accommodating large cargoes, both longstanding questions in the trafficking field. We anticipate that subsequent studies of the Sec24D/O-GlcNAc/MYOF axis and other novel Sec24D interactors will continue to advance our understanding of these processes in both health and disease.

## Experimental procedures

### Antibodies and reagents

The following antibodies were used: Rabbit anti-collagen I antiserum (LF-68) was a gift from L. Fisher, National Institute of Dental and Craniofacial Research; Rabbit monoclonal anti-Sec24D (D9M7L, #14687), Rabbit polyclonal anti-Sec23A (#8162), Rabbit MultiMab monoclonal mix anti-O-GlcNAc (#82332), Rabbit monoclonal anti-Calnexin (C5C9, #2679), Rabbit monoclonal anti-CD44 (E7K2Y, #37259), Rabbit monoclonal anti-IRS-1 (59G8, #2390), and Rabbit anti-HA (C29F4, #3724) from Cell Signaling Technology; Mouse anti-c-Myc (9E10, #626802), and Mouse anti-O-GlcNAc (RL2, #677902) from BioLegend; Mouse anti-O-GlcNAc (18B10.C7, #MA1-038) from ThermoFisher Scientific; Mouse anti-α-Tubulin (#T6074), Rabbit polyclonal anti-MIA3 (TANGO1) (#HPA055922), Rabbit polyclonal anti-Sec23IP (#HPA038403), and Rabbit polyclonal anti-SVIL (#HPA020138) from Sigma-Aldrich; Rat monoclonal anti-HA (3F10, #11867423001) from Roche; Rabbit polyclonal anti-Sec16A (#A300-648A), and Goat anti-myc tag antibody (#A190-104A) from Bethyl Laboratories; Mouse anti-Sec31A (32, #612351), and Mouse anti-GM130 (35, #610822) from BD Biosciences; Rabbit polyclonal anti-Sec13 (#A11613) from ABclonal; Rabbit polyclonal anti-Sec23B (#NBP2-20279) from Novus Biologicals; Rabbit monoclonal anti-Sec24D (EPR16089, #ab191566), Chicken anti-GFP (#ab13970), Rabbit monoclonal anti-LMAN1 (ERGIC53) antibody (EPR6979, #ab125006) and Rabbit monoclonal anti-MYOF (EPR18887, #ab178386) from Abcam; Mouse anti-Calnexin (2A2C6, #66903-1-Ig) from Proteintech; mouse anti-O-GlcNAc (CTD110.6, #sc-59623), and mouse anti-ERGIC53 (C-6, #sc-365158) from Santa Cruz Biotechnology; anti-procollagen I (PIP, #42024) from QED Bioscience; Goat horseradish peroxidase (HRP)-conjugated anti-mouse IgG (#1030-05), and Goat HRP-conjugated anti-rabbit IgG (#4030-05) from SouthernBiotech; Goat IRDye 800CW-conjugated anti-mouse IgG (#925-32210), and Goat IRDye 800CW-conjugated anti-rabbit IgG (#925-32211) from LI-COR; Goat Alexa Fluor 488-conjugated anti-mouse IgG (A-11001), Goat Alexa Fluor 488-conjugated anti-rabbit IgG (A-11008), Goat Alexa Fluor 488-conjugated anti-chicken IgY (A-11039), Goat Alexa Fluor 594-conjugated anti-mouse IgG (A-11005), Goat Alexa Fluor 594-conjugated anti-rabbit IgG (A-11012), Goat Alexa Fluor 647-conjugated anti-mouse IgG (A-21235), Goat Alexa Fluor 647-conjugated anti-rabbit IgG (A-32733), Goat Alexa Fluor Plus 647-conjugated anti-mouse IgG highly cross-adsorbed (A32728), Goat Alexa Fluor Plus 594-conjugated anti-rat IgG highly cross-adsorbed (A48264), Donkey Alexa Fluor 488-conjugated anti-mouse IgG (A-21202).

The following reagents and kits were used: Sodium ascorbate (Cat#A4034-100G) and Duolink In Situ PLA Probe anti-rabbit PLUS (Cat#DUO92002), Duolink In Situ PLA Probe anti-mouse MINUS (Cat#DUO92004), Duolink In Situ PLA Probe anti-goat MINUS (Cat#DUO92006), and Duolink In Situ Detection Reagents Red (Cat#DUO92008) from Sigma-Aldrich; DSP (Cat#22585), Lipofectamine RNAiMAX (Cat#13778150), and Lipofectamine 3000 (Cat#L3000015) from ThermoFisher Scientific; Biotin (#B4501-100MG). Thiamet-G (TG) from Cayman Chemical (#13237).

Per-acetylated 5S-GlcNAc (5SG) was synthesized by the Duke Small Molecule Synthesis Facility as described previously^115^.

The following siRNAs were purchased and used: siGENOME Human MYOF (26509) siRNA – SMARTpool (Dharmacon, M-013584-01-0005).

### Plasmid construction

Single guide RNAs (sgRNAs) for *SEC24D* knock-in (5’-TTATGAAAATATCATTCCAT-3’, designed with SYNTHEGO https://design.synthego.com/#/) and *MYOF* knock-out (#1: 5’-ACTGTCGGCCCTCAATCACT-3’) were subcloned into pSpCas9(BB)-2A-GFP (pX458, Addgene #48138) by the Duke Functional Genomics Facility. pX458 harboring sgRNA Sec24D#3 was reported previously^30^. For *SEC24D* knock-in repair templates, pAAVS1-3xFLAG-2xStrep (Addgene, #68375) was amplified with pAAVS-KI-F and pAAVS-KI-R primers. Gene blocks with left and right homology arms and myc-6xHis-tagged *SEC24D* sequence with a single mutation in the PAM sequence of a sgRNA for KI were designed and amplified with Repair-F and Repair-R primers. Amplified sequences of gene blocks were cloned into pAAVS1 with NEBuilder HiFi DNA Assembly Master Mix (New England BioLabs, #M5520AA) by mixing at 2:1 ratio at 50 °C for 1 h. To construct plasmids expressing O-GlcNAc deficient Sec24D mutants, pLenti6-myc-6xHis-hSec24D^30^ was mutated using Q5 Site-Directed Mutagenesis Kit (New England Biolabs, E0554) according to the manufacturer’s protocols with 9A-F and 9A-R primers and 13A-F and 13A-R primers for T9A and S13A, respectively. For MYOF expression, pcDNA3.1-MYOF-HA was gifted from Dr. William Sessa (Addgene, #22443)^116^. For the MYOF ΔC2A mutant, pcDNA3.1-MYOF-HA was amplified with MYOF-ΔC2A-F and pLenti-Gib-R2 primers and pLenti-Gib-F2 and MYOF-ΔC2A-R primers, respectively. Amplified sequences were assembled with NEBuilder HiFi DNA Assembly Master Mix (New England BioLabs, #M5520AA) by mixing them at 1:2 ratio at 50 °C for 1 h. For plasmids encoding the siRNA-resistant form of MYOF, a gene block with mutations against three siRNAs were designed and amplified with MYOF-si-resist-F1 and MYOF-si-resist-R2 primers. pcDNA3.1-MYOF-HA and pcDNA3.1-MYOF-ΔC2A-HA were amplified with pLenti-Gib-F2 and MYOF-si-resist-R1 primers, MYOF-si-resist-F2 and MYOF-si-resist-R3 primers, and MYOF-si-resist-F3 and pLenti-Gib-R2, respectively. Amplified sequences and an amplified gene block were assembled with NEBuilder HiFi DNA Assembly Master Mix (New England BioLabs, #M5520AA) by mixing at 1:1:1:1 ratio at 50 °C for 1 h. Primers and gene blocks used in this study are listed in Table S2.

### Cell culture

SW1353 cells were obtained from ATCC. Sec24D^WT^, Sec24D^T9A^, and Sec24D^S13A^ cell lines were cultured in Dulbecco’s modified Eagle’s medium (DMEM) supplemented with 10% fetal bovine serum at 37 °C under 5% v/v CO_2_.

### Inhibitor treatment

Cells were treated with 50 µM 5SG (OGT inhibitor) or 50 µM TG (OGA inhibitor) for 24 h at 37 °C.

### siRNA transfection

Cells at approximately 30% confluence, grown on appropriate dishes, were transfected with control or MYOF siRNAs using Lipofectamine RNAiMAX reagent, according to the manufacturer’s protocol, and were incubated for 48 hours for the experiments.

### Plasmid transfection

Cells at approximately 50% confluence, grown on appropriate dishes, were transfected with each plasmid using Lipofectamine 3000 reagent, in accordance with the manufacturer’s protocol, and were incubated for 24 hr. For rescue experiments with HA-MYOF, cells transfected with siRNAs for 24 hr were further transfected with each plasmid, as described above, and incubated for another 24 hr.

### Establishment of CRISPR knock-in (KI) and knockout (KO) cells

To generate KI cells expressing myc-His-tagged Sec24D, parental SW1353 cells grown on 15-cm dishes were co-transfected with pSpCas9(BB)-2A-GFP (pX458) harboring an sgRNA for KI and a pAAV-based repair template (Sec24D^WT^, Sec24D^T9A^, or Sec24D^S13A^ sequence) at 1:2 ratio. The next day, GFP-positive cells were collected by the Duke Cancer Institute Flow Cytometry Core Facility using a BD Diva fluorescence-activated cell sorter, followed by limiting dilution to isolate single cell clones. Homozygous KI cell lines for Sec24D^WT^ and Sec24D^S13A^ and a heterozygous KI cell line for Sec24D^T9A^ were obtained. To obtain homozygous Sec24D^T9A^ mutant cells, the heterozygous KI clone was co-transfected with pX458 with sgRNA for Sec24D#3, reported previously^30^, and a pAAV-based repair template with Sec24D^T9A^ sequence at 1:2 ratio. GFP-positive cells were collected and subjected to limiting dilution. A homozygous KI cell line for Sec24D^T9A^ was obtained.

For MYOF-KO cells, parental SW1353 cells grown on 6-cm dishes were transfected with plasmids with sgRNAs targeting MYOF. The next day, GFP-positive cells were sorted as described above, followed by limiting dilution. A MYOF-KO clone derived from cells transfected with sgRNA#1 was validated and used for experiments.

### Genotyping of KI cells

Parental and KI cells were grown on 12-well plates, and genomic DNA was extracted in 100 µl GNT buffer containing 1 µl proteinase K (Thermo Fisher, EO0492) by incubating at 55 °C for more than 1.5 h. Proteinase K was heat-inactivated at 95 °C for 5 min. PCR was carried out with 24D-KI-geno-F and 24D-KI-geno-R primers. For sequencing, PCR was conducted using hSec24D-KI-seq-F and hSec24D-KI-seq-R primers.

### Sample preparation for IB

Cells were washed with phosphate-buffered saline (PBS) twice and collected using cell scrapers, followed by precipitation by centrifugation at 1,000 × g for 3 min. Cell pellets were lysed in 8 M urea in PBS by sonication, followed by centrifugation at 22,640 × g for 15 min at 4 °C. Supernatants were recovered, and protein concentration was measured by BCA assay. Samples were mixed with 5× SDS loading buffer and incubated at 95 °C for 5 min.

### IB

Equal amounts of proteins were loaded in each well and separated by Tris-glycine SDS-PAGE. For enhanced chemiluminescence (ECL), SDS-PAGE gels were electroblotted onto PVDF membranes (88515, ThermoFisher). The membranes were blocked with blocking buffer [2.5% BSA in Tris-buffered saline with 0.1% Tween-20 (TBS-T)]for 30 min, followed by incubation overnight at 4 °C with primary antibodies diluted in blocking buffer. After washing with TBS-T three times, membranes were incubated with secondary antibodies conjugated with HRP diluted 1:10,000 in blocking buffer for 1 h at room temperature. Membrane were washed with TBS-T three times and signal was detected via ECL (Genesee Scientific 20-300B) and photographic film (LabScientific, XAR ALF 2025). For quantitative fluorescent IBs, gels were electroblotted onto nitrocellulose membranes (162-115, BioRad). After blocking, incubation with primary antibodies, and washing with the protocol described above, membranes were incubated with secondary antibodies conjugated with IRDye800 diluted 1:20,000 in blocking buffer for 1 h at room temperature. Membranes were washed with TBS-T three times, followed by washing with TBS one time, and signals were detected using a LI-COR Odyssey DLx Imaging System. Images were quantified with Fiji. The following primary antibodies were used: 18B10 (1:1,000); CTD110.6 (1:200); myc (mouse, 1:1,000); α-Tubulin (1:50,000); Sec24D (from Abcam, 1:1,000); Sec24A (1:1,000); Sec24B (1:20,000); Sec24C (1:5,000); COL1A1 (rabbit, 1:5,000); MultiMab (1:1,000); Sec23A (1:2,000); Sec31A (1:500); Sec23B (1:1,000); Sec13 (1:10,000); TANGO1 (1:500); Sec16A (1:500); Sec23IP (1:500); MYOF (1:1,000); IRS1 (1,1000); SVIL (1:500); pan-14-3-3 (1:1,000); Calnexin (rabbit, 1:1,000); CD44 (1:1,000); HA (rabbit, 1:2,000).

### Collagen accumulation assay

Cells grown on 18-mm coverslips in 24-well plates were treated with 50 µg/ml ascorbate at 37°C for 5 h to allow collagen production and secretion. Cells were then washed with cold PBS twice and fixed with 4% paraformaldehyde (PFA) at room temperature for 15 min, followed by washing with cold PBS three times. Cells were permeabilized with PBS-T (PBS containing 0.3% Triton X-100) for 10 min at room temperature and incubated with blocking solution, primary antibodies, and secondary antibodies as described below.

For IB experiments, cells grown on 6-cm dishes were transfected with empty vector or plasmids encoding Sec24D^WT^, Sec24D^T9A^, or Sec24D^S13A^. The next day, cells were treated with 50 µg/ml ascorbate at 37 °C for 5 h, followed by sample preparation for IB as described above.

### Collagen transport assay

For kinetic collagen transport assays, cells were grown on 18-mm coverslips in 24-well plates and pre-incubated at 40 °C supplemented with 20 mM HEPES-NaOH (pH 7.4) for 3 h. Soon after preincubation, cells for 0 min samples were washed with cold PBS twice and fixed with 4% PFA at 32°C for 15 min. Culture medium for the remainder of the samples was replaced with pre-warmed medium supplemented with 20 mM HEPES-NaOH (pH 7.4), 100 µg/ml cycloheximide, and 50 µg/ml freshly dissolved ascorbate, and cells were incubated at 32 °C for 10 or 15 min. After transport, cells were washed with cold PBS twice and fixed with 4% PFA at 32 °C for 15 min. After fixation, cells were washed with PBS three times, followed by permeabilization with PBS-T for 10 min at room temperature. After permeabilization, cells were incubated with blocking solution, primary antibodies, and secondary antibodies as described below.

For quantifying O-GlcNAc levels of Sec24D in response to collagen transport by IP, cells grown on 15-cm dishes were pre-incubated at 40 °C, as above. Cells were placed on ice and washed with cold PBS twice (for 0 min). Culture medium for the remainder of the samples was replaced with pre-warmed medium supplemented with 20 mM HEPES-NaOH (pH 7.4), 100 µg/ml cycloheximide, and 50 µg/ml ascorbate, and cells were incubated at 32 °C for 5 min. Cells were then quickly placed on ice and washed with cold PBS twice. Cells were harvested by scraping on ice and centrifuged at 500 × g for 5 min at 4°C. After discarding the supernatants, cells were subjected to tandem purification as described below.

### E-Cadherin RUSH transport assay

Cells grown on 18-mm coverslips in 24-well plates were transfected with a plasmid encoding the E-Cadherin RUSH probe. The next day, cells were washed with cold PBS twice and fixed with 4% PFA at 37 °C for 15 min (for 0 min samples). Culture medium for the remainder of the samples was replaced with pre-warmed medium supplemented with 100 µg/ml cycloheximide, and 50 µM biotin, and cells were incubated at 37 °C for 20 or 40 min. After transport, cells were washed with cold PBS twice and fixed with 4% PFA at 37 °C for 15 min. After fixation, cells were permeabilized, incubated with blocking solution, primary antibodies, and secondary antibodies as described below.

### IF

Cells were grown on 18-mm coverslips in 24-well plates. After washing with PBS twice, cells were fixed with cold methanol for 15 min at -20 °C. Cells were washed with PBS three times, followed by incubation with blocking solution (PBS containing 5% goat serum, 0.3% Triton X-100, and 0.05% NaN_3_) for 30 min at room temperature and incubated with primary antibodies diluted in blocking solution at 4 °C overnight. After washing with PBS three times, cells were incubated with secondary antibodies conjugated with Alexa dyes diluted 1:1,000 in blocking solution for 1 h at room temperature. After washing with PBS-T three times, coverslips were mounted on slides with ProLong Diamond anti-fade mounting medium. Images were acquired by Zeiss 780 with a 40×/1.4 NA Oil Plan-Apochromat DIC, (UV) VIS-IR or a 63×/1.4 NA Oil Plan-Apochromat DIC oil immersion objective lens for collagen and E-Cadherin RUSH transport assays or other experiments, respectively. Z-stack images were obtained. Images were processed using FIJI Image J software.

To detect transfected proteins, cells grown on 18-mm coverslips in 24-well plates were transfected with empty vector or plasmids encoding HA-MYOF (FL or ΔC2A). After washing with PBS twice, cells were fixed with 4% PFA for 15 min at room temperature. After fixation, cells were washed with PBS three times, followed by permeabilization with PBS-T for 10 min at room temperature. After permeabilization, cells were incubated with blocking solution, primary antibodies, and secondary antibodies as described above.

The following primary antibodies were used: myc (mouse, 1:1,000); COL1A1 (rabbit, 1:2,000; mouse, 1:2,000, 1 µg/ml); Calnexin (mouse, 1:500); GM130 (1:500); GFP (1:2,000); Sec24D (CST, 1:300); Sec16A (1:300); Sec31A (1:300); ERGIC53 (rabbit, 1:300); HA (rat, 1:2,000, 0.05 µg/ml).

### PLA

PLA assays were carried out according to the manufacturer’s protocol. Cells were grown on 12-well chamber slides (ibidi, Cat#81201) and fixed in cold methanol at -20 °C for 15 min. After washing with PBS three times, cells were incubated in Blocking Solution (supplied in kit) at 37 °C for 30 min. Subsequently, cells were incubated with primary antibodies [mouse anti-myc (1:1,000) and rabbit anti-O-GlcNAc (MultiMab, 1:500), rabbit anti-MYOF (1:100), or rabbit anti-Collagen I (1:500)] dissolved in Antibody Diluent (supplied in kit) at 4 °C overnight. After washing with 1 × Wash buffer A (supplied in wash buffer kit, Sigma-Aldrich, Cat#DUO82049) for 2 min twice, cells were incubated with PLA probes for mouse and rabbit diluted 1:5 in Antibody Diluent at 37 °C for 60 min. After washing with 1× Wash buffer A for 2 min twice, cells were incubated with Ligation-Ligase solution at 37 °C for 30 min. After washing with 1 × Wash buffer A for 2 min twice, cells were incubated with Amplification-Polymerase solution at 37 °C for 100 min. After washing with 1 × Wash buffer B (supplied in the wash buffer kit) for 10 min twice and with 0.01 × Wash buffer B for 1 min once, the silicon separator was removed and the slide was mounted on cover glass.

For co-staining with Sec31A or ERGIC53, cells fixed with cold methanol at -20 °C for 15 min were incubated in 1% BSA solution dissolved in supplied Blocking Solution at 37 °C for 30 min and were incubated with primary antibodies [goat anti-myc (1:5,000) and rabbit anti-O-GlcNAc (MultiMab, 1:500) or rabbit anti-MYOF (1:100)] dissolved in Antibody Diluent at 4 °C overnight. The rest of the PLA assay was performed as above. After the second wash with 1 × Wash buffer B, cells were further stained with primary antibody [mouse anti-Sec31A (1:100) or mouse anti-ERGIC53 (1:100)] for 1 h at room temperature. Cells were washed with PBS for 5 min three times and stained with secondary antibody conjugated with Alexa Fluor 488 for 1 h at room temperature. After washing with PBS-T for 5 min three times, the silicon separator was removed and the slide was mounted on cover glass.

For combined PLA and collagen transport assay, cells were fixed with 4% PFA at 37 °C for 15 min, followed by washing with PBS for 5 min three times. Cells were then permeabilized with PBS-T for 10 min at room temperature. Subsequently, cells were incubated in Blocking Solution at 37 °C for 30 min, followed by incubation with primary antibodies [mouse anti-myc (1:1,000) and rabbit anti-O-GlcNAc (MultiMab, 1:500)] dissolved in Antibody Diluent at 4 °C overnight. The rest of the PLA assay was performed as above, and the slide was mounted on cover glass.

### Quantification of IF data

All images were obtained as z-stacks and processed with Fiji. Before analyzing, all images were projected with Max Intensity and subjected to background subtraction with a rolling-ball radius with 50.0 pixels. For collagen accumulation experiments, mean fluorescence intensities (MFIs) of collagen were measured. For collagen transport experiments, Pearson’s correlation coefficient (PCC) between COL1A1 and GM130 was calculated with the Colo2 plug-in. Cells with PCC ≥ 0.45 were considered Golgi-localized, and the percentage was calculated by dividing by total cell number. For E-Cadherin RUSH assays, FIs of GFP were measured to obtain total FIs. Images of Golgi-localized GFP were created by imageCalculator with the “AND create” algorithm with GFP and GM130 images to extract regions of overlap, and FIs of Golgi-localized GFP were measured in them. Mander’s overlapping coefficient (MOC) was calculated with Colo2 plug-in in images produced with appropriate thresholding methods. Costes test was conducted on each cell to determine whether the obtained MOCs were significantly higher than those from random overlap in the same image. A Costes *p* value ≥ 0.95 was deemed to indicate that the corresponding MOCs are not due to random overlap. MFIs of Sec31A in puncta were measured in images produced with the appropriate thresholding method with Analyze Particles plug-in and default settings. To quantify particle numbers and size of COL1A1 in MYOF-KD cells, images at 15 min after collagen release were used. Images of COL1A1 staining were subtracted with GM130 images by imageCalculator using the “Subtract create” algorithm to exclude Golgi-localized COL1A1 from the analyses. The generated images were analyzed with Analyze Particles with size = 0.70 – ∞ and circularity = 0.75 – 1.00.

### Quantitative reverse-transcription PCR (qRT-PCR)

Cells grown on 6-well plates were harvested after washing with PBS, and total RNA was extracted with the RNeasy Mini Kit according to the manufacturer’s protocol (Qiagen, #74104). RNA concentration was measured by Nanodrop 2000 Spectrophotometer (ThermoFisher, ND-2000). cDNAs were synthesized from 1 µg total RNA using SuperScript IV reverse transcriptase (ThermoFisher, #18090050) with 50 µM random hexamers according to the manufacturer’s protocol. qRT-PCR was conducted with Power SYBR Green PCR Master Mix (Applied Biosystems, #4367659) according to the manufacturer’s protocol using the following primer sets: hCOL1A1, hCOL1A1-RT-F and hCOL1A1-RT-R; hACTB, b-actin-RT-F and b-actin-RT-R. Relative mRNA levels of COL1A1 compensated by mRNA levels of beta-actin were calculated by ΔΔCT method.

### Fish maintenance and breeding

The zebrafish *bulldog* allele m494 (*bul^m^*^494^) was raised under standard laboratory conditions at 28.5 °C with a constant photoperiod (14 h light: 10 h dark) as previously described^15^ by the Institutional Animal Care and Use Committee at Vanderbilt University Medical Center (protocol number M1700020). Zebrafish larvae were imaged at indicated developmental stages [days post-fertilization (dpf)].

### Zebrafish microinjections

The human *SEC24D*^WT^ gene was cloned into the pDONR221 vector using Gateway BP Clonase II enzyme mix (11789020, ThermoFisher Scientific). Subsequently, the multistep Gateway Recombineering system (p5E_1.7kb Col2a1a promoter, p3E_v2a-EGFP, and pDestTol2pA2) and Gateway LR Clonase II enzyme (11791020, ThermoFisher Scientific) were used to create the SEC24D fusion construct with a self-cleaving viral 2a peptide-tagged EGFP (v2a-EGFP) under the tissue-specific Col2a1a promoter. SEC24D O-GlcNAc sites were mutated using the Q5 site-directed mutagenesis kit (E0554S, NEB) with the following primers: ACA-GCA Threonine to Alanine (T9A) Fwd: TTACGTGGCTgCACCTCCGTA, Rev: CCTTGTTGACTCATAGCCTG; TCT-GCT, Serine to Alanine (S13A) Fwd: ACCTCCGTATgCTCAGCCTCA, Rev: GTAGCCACGTAACCTTGTTG.

Zebrafish larvae at the single-cell stage were collected from *bul^m^*^494^*/AB* heterozygous crosses, injected with a combination of 100 pg of medaka transposase mRNA and 10 pg of SEC24D vector DNA, and grown for 4 days in larvae medium^31^.

### Zebrafish cryosectioning and immunohistochemistry

Cryosectioning and immunohistochemistry were performed essentially as described^31^. Briefly, four dpf larvae were fixed with 4% PFA at 4 °C overnight, then washed and transferred to 30% sucrose in PBS. Larvae were embedded in Cryomatrix (6769006, ThermoFisher Scientific) and frozen at −80 °C for 15 min. Then, 14-μm-thick sections were cut with a cryostat (Leica CM1900) and collected onto Fisher brand Superfrost Plus slides (12-550-15, Thermo Fisher Scientific). Tissue sections were dried and rehydrated in PBS before staining. Samples were further processed with 20 μg/mL proteinase K (P50220 RPI) at room temperature for 5 min for antigen retrieval, and larvae were permeabilized with 0.5% Triton X-100 in PBS for 10 min at room temperature^117^. Tissue Samples were blocked [2% goat serum (Sigma, G9023) and 2 mg/mL BSA (Sigma, A8022)] for 30 min at room temperature, followed by overnight primary antibody incubation at 4 °C, using collagen II antibody (DSHB, 116B3; 1:250) and GFP antibody (Vanderbilt University Antibody and Protein Resource; 1:250). Samples were incubated with secondary antibodies 1:500 [mouse AlexaFluor-555 (Life Technologies, A-21422) and chicken AlexaFluor-488 (Life Technologies, A-11039)] for one hour and 1:4000 DAPI (D1306 Invitrogen) for 15 min at room temperature. Slides were mounted in Prolong Gold (P36930, Thermo Fisher Scientific).

### Zebrafish imaging and quantification

Samples were imaged using an AxioImager Z1 equipped with an Apotome and an EC Plan Neofluar 100X/1.30 Oil objective and ORCA-Flash4.0 Digital CMOS Camera. Images were analyzed in FIJI^118^. The percent collagen area in the cytosol was calculated by the following formula: [collagen-positive intracellular area/(cytosol area − nucleus area marked by DAPI)] × 100. ImageJ was used to measure intracellular areas^31^.

### Chemical crosslinking with DSP

Cells grown on 15-cm dishes were washed with cold PBS twice, followed by incubation with PBS containing 0.1 mM DSP at 4 °C for 2 h. After discarding PBS containing DSP, excess DSP was quenched with STOP solution [20 mM Tris-HCl (pH 7.4) in PBS] by incubating at room temperature for 15 min. Cells were then harvested by scraping, followed by tandem purification as described below.

### Tandem purification

For silver staining and MS analysis, the following protocol was used to purify Sec24D and associated proteins. Cells crosslinked with DSP were lysed by sonication in lysis buffer containing 20 mM Tris HCl (pH 7.4), 150 mM NaCl, 1% Triton X-100, 0.1% SDS, and protease inhibitor cocktail (Sigma-Aldrich, P8340, 1:100) followed by centrifugation at 22,640 x g for 15 min at 4 °C. Protein concentration was measured by BCA and normalized. Samples were incubated with anti-myc antibody (2 µg antibody per 1 mg total protein) at 4 °C for 1 h, followed by incubation with 30 µl of protein A/G UltraLink resin (ThermoFisher, 53133) at 4 °C overnight. After washing beads three times with lysis buffer, bound proteins were eluted with 500 µl of Ni-wash buffer (300 mM NaCl, 1% Triton X-100, and 10 mM imidazole in 8 M urea/PBS) twice, each for 15 min, with rotation at room temperature. The two eluates were combined and rotated with 50 μL of HisPur Ni-NTA resin (ThermoFisher, 88223) for 2 h at room temperature. Resin was then washed with Ni-wash buffer three times, and eluted with 40 µl of elution buffer (250 mM imidazole in 8 M urea/PBS) for 20 min with rotation at room temperature. For mass spectrometry, 30 µl of eluates were reduced by incubating in 20 mM DTT at room temperature for 30 min. The samples were stored at -80 °C until MS analysis. For silver staining, 40 µl of eluates were mixed with 5× SDS loading buffer and incubated at 95 °C for 5 min.

For O-GlcNAc analysis, cells were lysed by sonication in lysis buffer containing 20 mM Tris HCl (pH 7.4), 150 mM NaCl, 1% Triton X-100, 0.1% SDS, protease inhibitor cocktail, 50 µM UDP (Sigma-Aldrich, 94330; OGT inhibitor), and 5 µM PUGNAc (Cayman Chemical, 17151; OGA inhibitor). Tandem-purification was conducted as described above, and captured proteins on Ni-NTA resin were eluted with 80 µl of elution buffer (250 mM imidazole in 8 M urea/PBS) for 20 min with rotation at room temperature. The eluates were mixed with 5× SDS loading buffer and incubated at 95°C for 5 min.

For co-IP experiments without DSP, cells were lysed by sonication in lysis buffer containing 20 mM Tris HCl (pH 7.4), 150 mM NaCl, 1% Triton X-100, 1 mM EDTA, 10% glycerol, protease inhibitor cocktail, 50 µM UDP, and 5 µM PUGNAc. The samples with equal amount of proteins were pre-cleared by incubating with 30 µl of protein A/G resin equilibrated with the same lysis buffer for 1 h at 4 °C. The resin was precipitated by centrifugation at 1,000 x g for 1 min, and the supernatants were collected for IP. The supernatants were incubated with anti-myc antibody (add 2 µg antibody per 1 mg total protein) for 1 h at 4 °C, followed by incubation with 30 µl of protein A/G resin at 4 °C overnight. After washing beads three times with 1 ml of IP wash buffer containing 20 mM Tris-HCl pH 7.4, 150 mM NaCl, 1% Triton-X 100, 10% glycerol, bound proteins were eluted with 500 µl of Ni-binding buffer [20 mM Tris-HCl pH 7.4, 150 mM NaCl, 1% Triton-X 100, 10% glycerol, 10 mM imidazole, 0.1 mg/ml 2 x Myc-peptide (ChromoTek, 2yp-1), protease inhibitor cocktail, 50 µM UDP (OGT inhibitor), and 5 µM PUGNAc (OGA/hexosaminidase inhibitor)] by incubating them for 1 h at room temperature. After spinning down the beads, supernatants were collected and incubated with 50 µl of Ni-NTA beads equilibrated by Ni-binding buffer for 2 h at room temperature. After washing beads with 1 ml of Ni-binding buffer three times, proteins were eluted with 80 µl of Ni-elution buffer containing 20 mM Tris-HCl pH 7.4, 150 mM NaCl, 1% Triton X-100, and 250 mM imidazole by incubating for 20 min at room temperature. The eluates were mixed with 5× SDS loading buffer and incubated at 95 °C for 5 min.

### Silver staining

Tandem-purified protein samples were separated by Tris-glycine SDS-PAGE with a precast 4-15% gradient gel (BioRad, 4561086). The gel was quickly washed with ultra-pure water three times with shaking and fixed with 100 ml of 40% ethanol/10% acetic acid with gentle shaking at room temperature for 1 hr. After washing in 30% ethanol for 10 min, the gel was sensitized, stained, and developed with SilverQuest Silver Staining Kit (ThermoFisher Scientific, LC6070) according to the basic staining method of the manufacture’s protocol.

### Sample preparation for mass spectrometry

Samples were spiked with 1 or 2 pmol bovine casein as an internal quality control standard, brought to 5% SDS and reduced for 15 min at 80 °C, alkylated with 20 mM iodoacetamide for 30 min at room temperature, then supplemented with a final concentration of 1.2% phosphoric acid and 375 µL of S-Trap (Protifi) binding buffer (90% MeOH/100mM TEAB) (Table S3). Proteins were trapped on the S-Trap micro cartridge, digested using 20 ng/µL sequencing grade trypsin (Promega) for 1 hr at 47 °C, and eluted using 50 mM TEAB, followed by 0.2% formic acid (FA), and lastly using 50% acetonitrile (ACN)/0.2% FA. All samples were then lyophilized to dryness. Samples were resuspended in 250 ng/µL of 1% trifluoroacetic acid (TFA)/2% ACN with 12.5 fmol/µL of yeast ADH. A study pool QC (SPQC) was created by combining equal volumes of each sample.

### Liquid chromatography (LC)-MS/MS analysis

Quantitative LC/MS/MS was performed on 2 µL of each sample, using a Vanquish Neo UPLC system (ThermoFisher) coupled to a Thermo Orbitrap Astral high-resolution accurate mass tandem mass spectrometer (ThermoFisher). Briefly, the sample was first trapped on a Symmetry C18 20 mm × 180 µm trapping column (5 μl/min at 99.9/0.1 v/v water/ACN), after which analytical separation was performed using a 1.5 µm EvoSep 150um ID x 8cm performance (EvoSep) column with a 30 min gradient of 5 to 30% ACN with 0.1% FA at a flow rate of 500 nL/min with a column temperature of 50 °C. Data collection on the Orbitrap Astral mass spectrometer was performed in a data-independent acquisition (DIA) mode of acquisition with a r=240,000 (@ m/z 200) full MS scan from m/z 380-980 with a target AGC value of 4e5 ions. Fixed DIA windows of 4 m/z from m/z 380 to 980 DIA MS/MS scans were acquired in the Astral with a target AGC value of 5e4 and max fill time of 6 ms. HCD collision energy setting of 27% was used for all MS2 scans. The total analysis cycle time for each sample injection was approximately 35 min.

### Quantitative MS data analysis

Following 15 total UPLC-MS/MS analyses (Table S3), data were imported into Spectronaut (Biognosis), and individual LCMS data files were aligned based on the accurate mass and retention time of detected precursor and fragment ions. Relative peptide abundance was measured based on MS2 fragment ions of selected ion chromatograms of the aligned features across all runs. The MS/MS data was searched against the SwissProt *Human* database (downloaded in 2022), a common contaminant/spiked protein database (bovine albumin, bovine casein, yeast ADH, etc.), and an equal number of reversed-sequence “decoys” for false discovery rate determination. A library-free approach within Spectronaut was used to perform the database searches. Database search parameters included fixed modification on Cys (carbamidomethyl) with variable modification on Met (oxidation). Full trypsin enzyme rules were used along with 10 ppm mass tolerances on precursor ions and 20 ppm on product ion. Spectral annotation was set at a maximum 1% peptide false discovery rate based on q-value calculations. Peptide homology was addressed using razor rules, in which a peptide matched to multiple different proteins was exclusively assigned to the protein with more identified peptides. Protein homology was addressed by grouping proteins that had the same set of peptides to account for their identification. A master protein within a group was assigned based on percent coverage.

Raw intensity values for each precursor (individual charge states) identified peptide are presented in Table S3. These values are directly from Spectronaut, which applies its own criteria for peak detection. If it does not meet those criteria, no value (i.e. a blank cell) is given. The most likely reason for this is if the peak is below the limit of detection (low abundance). Next, the following imputation strategy was applied to missing values. First, peptides with the sequence but different precursor charges state were summed to give a single peptide value. Then, if less than half of the values are missing in a biological group, values are imputed with an intensity derived from a normal distribution of all values defined by measured values within the same intensity range (20 bins). If greater than half the values are missing for a peptide in a group and a peptide intensity is > 5e6, then it was concluded that peptide was misaligned and its measured intensity is set to 0. All remaining missing values are imputed with the lowest 2% of all detected values (Table S3). Prior to normalization, a filter was applied such that a peptide was removed if it was not measured at least twice across all samples and in at least 50% of the replicates in any one single group. After that filter, samples were total intensity-normalized (i.e., total intensity of all peptides for a sample are summed ad then normalized across all samples). These data were then subjected to a trimmed-mean normalization, in which the top and bottom 10 percent of the signals were excluded and the average of the remaining values was used to normalize across all samples. Lastly, all peptides belonging to the same protein were summed into a single intensity (Table S3). These normalized protein level intensities were used for the remainder of the analysis.

### Sequence alignment of Sec24D

Sequence alignment was conducted with Clustal Omega (https://www.ebi.ac.uk/jdispatcher/msa/clustalo) and visualized with Boxshade (https://junli.netlify.app/apps/boxshade/). Amino acids sequences were obtained from NCBI: human Sec24D (NP_055637.2); mouse Sec24D (NP_081411.2); rat Sec24D (NP_001296382.1); zebrafish Sec24D (NP_001171403.2).

### GO enrichment analysis

GO enrichment analysis was conducted with Metascape (https://metascape.org/gp/index.html#/main/step1). Three hundred sixty genes specifically identified in Sec24D^WT^ cells were analyzed.

### Generation of interaction maps

Interaction maps of proteins were generated with STRING (https://string-db.org). Proteins with more than two identified peptides were considered strong candidates for this analysis.

### Prediction of subcellular localization on endogenous MYOF

Subcellular localization of endogenous MYOF was predicted with The HeLa Spatial Proteome (http://mapofthecell.biochem.mpg.de) and a deposited dataset reported previously^78^.

### Statistics

Statistical analyses were performed using GraphPad Prism 9 or 10 (GraphPad Software, Inc.).

### Resource availability

#### Lead contact

Requests for further information and resources should be directed to and will be fulfilled by the lead contact, Michael Boyce.

#### Materials availability

All unique/stable reagents generated in this study are available from the lead contact with a complete materials transfer agreement.

#### Data and code availability

Proteomics data of interactome analyses in parental, Sec24D^WT^, Sec24D^T9A^, and Sec24D^S13A^ SW1353 cells have been deposited to the ProteomeXchange Consortium via PRIDE partner repository and are available via ProteomeXchange with identifier PXD058582 (reviewer can access the dataset by logging in to the PRIDE website using the following details: Project accession: PXD058582; Token: B4eeSUJLPZKb). All other data reported in this paper will be shared by the lead contact upon request. This paper does not report original code.

## Acknowledgments

We thank Dr. So Young Kim at the Duke Functional Genomics Core for construction of plasmids encoding sgRNAs, Dr. Bin Li at the Duke Cancer Institute Flow Cytometry Shared Resource for cell sorting, Tricia Ho and Greg Waitt at the Duke Proteomics and Metabolomics Shared Resource for MS data acquisition, and Cory L. Guthrie for excellent zebrafish care at VUMC. We also thank members of the Boyce lab for constructive discussions. TH was supported by The Osamu Hayaishi Memorial Scholarship for Study Abroad from the Japanese Biochemical Society. This work was supported by grant 5R01GM117473 from the National Institute of General Medical Sciences to M.B. and R01MH113362 from the National Institute of Mental Health to E.W.K.

## Author contributions

Conceptualization: TH, MB; Formal analysis: TH, DC, ES; Investigation: TH, DC, ES; Resources: TH, DC, BB, EWK; Writing – Original Draft: TH; Writing – Review & Editing: TH, DC, EWK, MB; Visualization: TH; Supervision: MB; Project Administration: TH, MB; Funding Acquisition: MB, EWK.

## Declaration of interest

The authors declare that they have no conflicts of interest regarding the contents of this article.

**Table S1.** Lists of proteins identified in MS proteomics analyses of Sec24D interactomes, related to Fig. 3.

**Table S2.** Oligonucleotides used in this study.

**Tables S3.** Source data files from proteomics analyses, related to Fig. 3.

---

**Figure S1.**
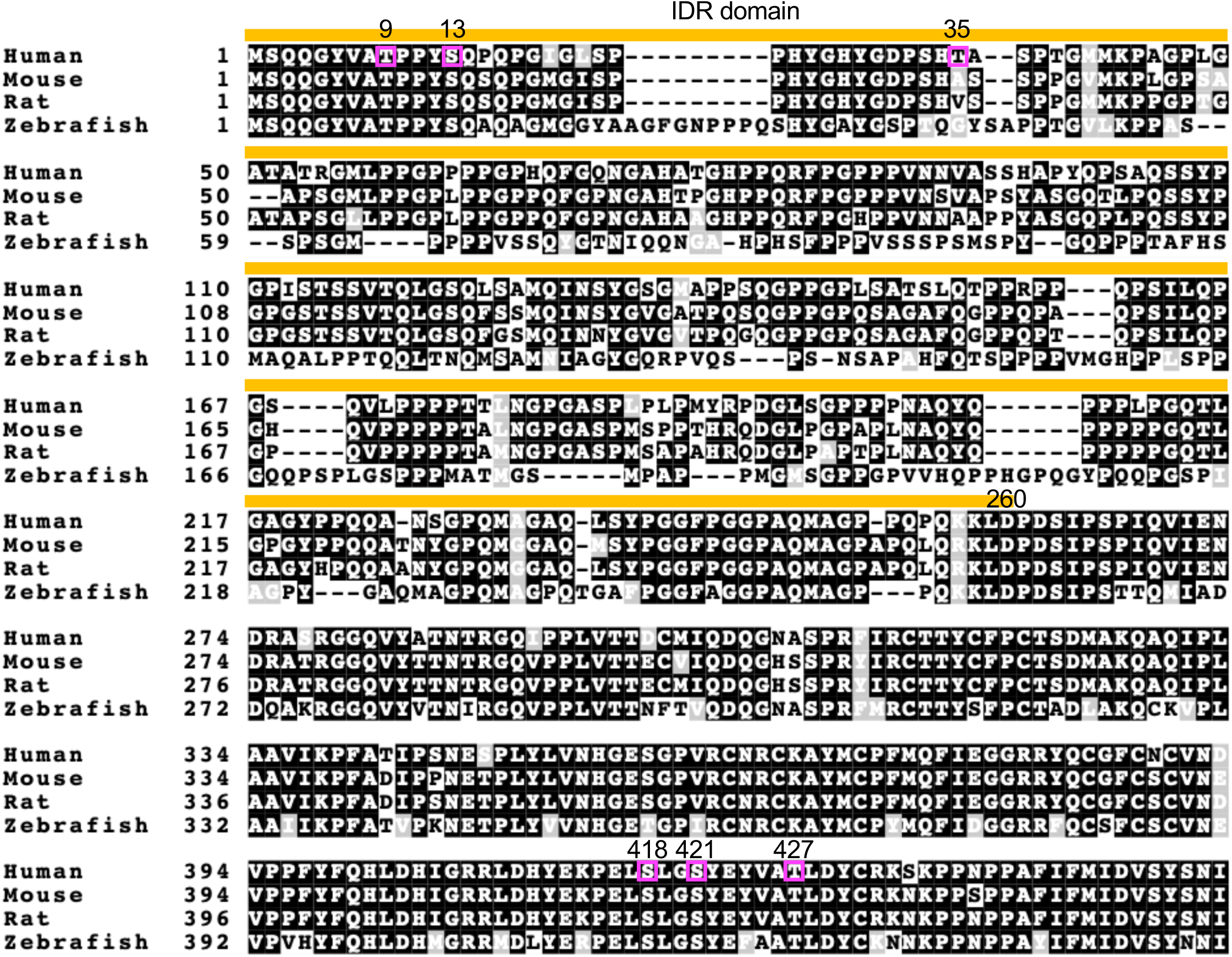
Alignment of vertebrate Sec24D sequences, related to Fig. 1. Amino acid sequences of the N-terminal regions of Sec24D orthologs from human, mouse, rat, and zebrafish were aligned. The IDR domain is highlighted with an orange line. O-GlcNAc sites previously identified in human Sec24D^30^ are highlighted in magenta.

**Figure S2.**
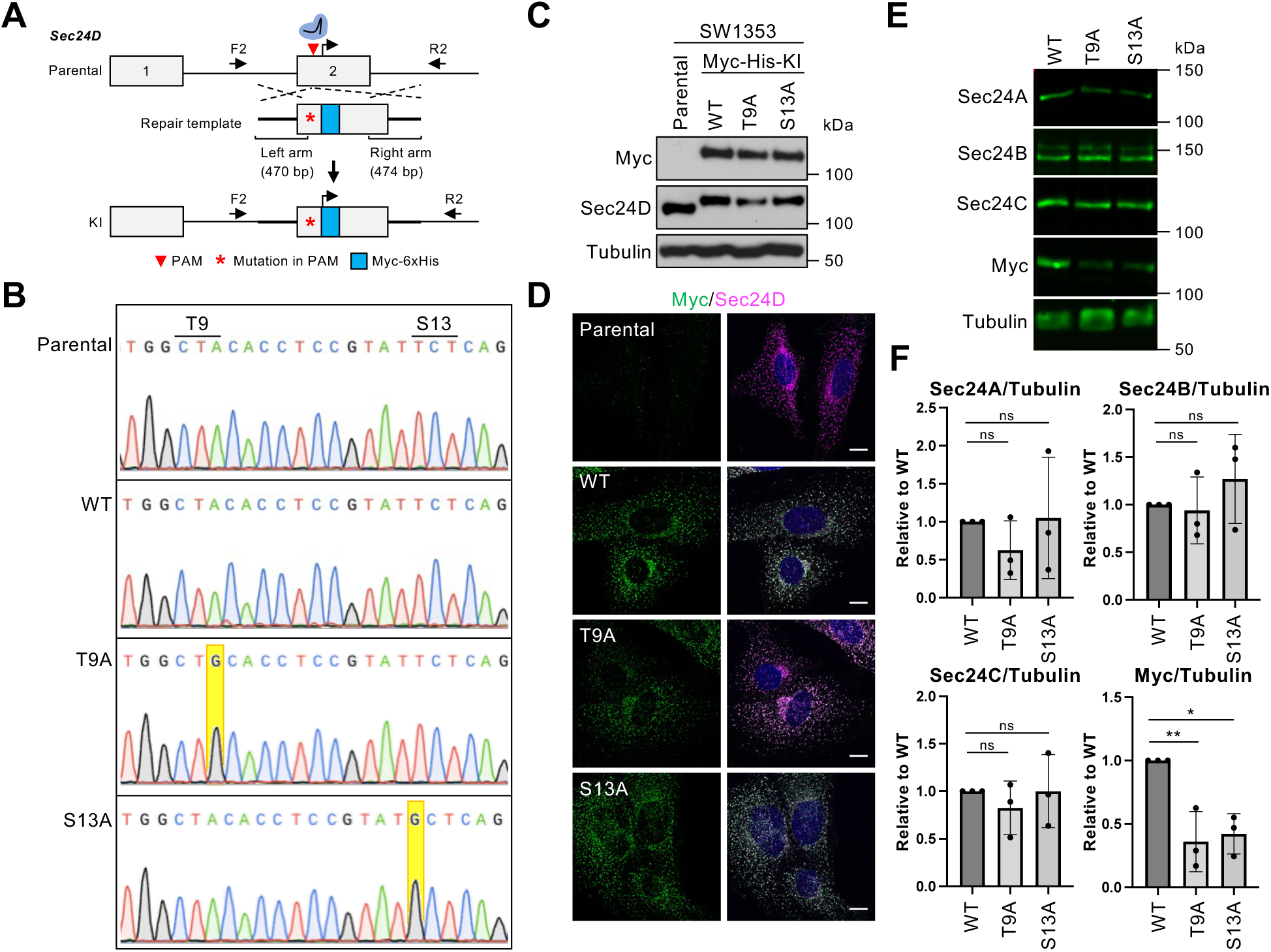
Generation of myc-6xHis-tagged *Sec24D* knock-in (KI) cells, related to Fig. 1. A. CRISPR strategy to generate *Sec24D* knock-in cells. F2 and R2 primers were used for Sanger sequencing of the entire regions of interest. B. Sequences of parental SW1353 and KI cells. Correct mutations in KI cells are highlighted in yellow. C. Parental SW1353, Sec24D^WT^, Sec24D^T9A^, and Sec24D^S13A^ cells were analyzed by IB. D. Parental SW1353, Sec24D^WT^, Sec24D^T9A^, and Sec24D^S13A^ cells were analyzed by IF. Scale bars: 10 µm. E. Sec24D^WT^, Sec24D^T9A^, and Sec24D^S13A^ cells were analyzed for Sec24 paralog expression levels by IB. F. Quantification of experiments depicted in E. The ratio of Sec24 paralog to tubulin (loading control) signals was normalized to Sec24D^WT^ (n = 3). Data were analyzed by one-way ANOVA with *post hoc* Tukey test. * p < 0.05; ** p < 0.01; ns: not significant.

**Figure S3.**
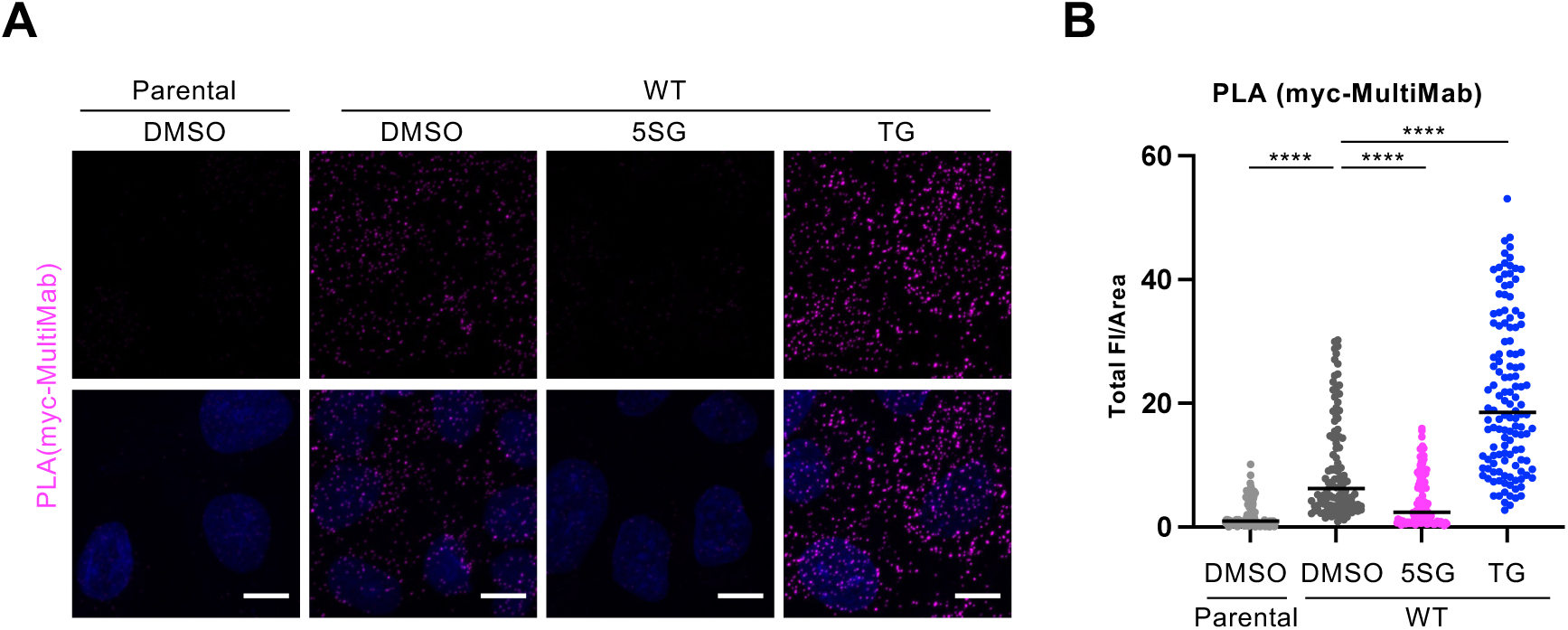
Validation of PLA for O-GlcNAcylated Sec24D, related to Fig. 1. A. Parental SW1353, Sec24D^WT^, Sec24D^T9A^, and Sec24D^S13A^ cells were treated with 50 µM 5SG or 50 µM TG for 24 hr, and O-GlcNAcylated Sec24D was detected by PLA (myc and MultiMab O-GlcNAc antibodies). B. Quantification of experiments depicted in A. Total FI of PLA signal was normalized to cell area (n = 3). Data were analyzed by two-way ANOVA with *post hoc* Tukey test. **** *p* < 0.0001.

**Figure S4.**
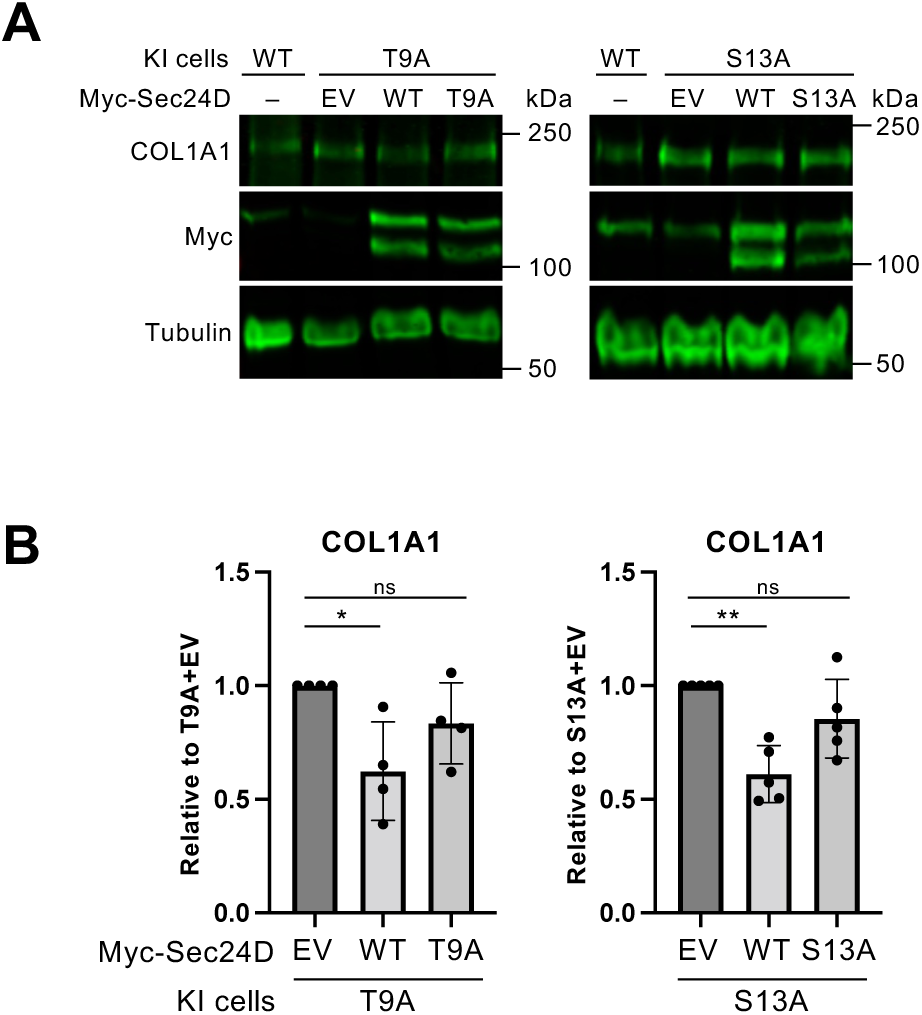
Overexpression of Sec24D^WT^, but not Sec24D^T9A^ or Sec24D^S13A^, rescues collagen transport defects in Sec24D^T9A^ or Sec24D^S13A^ cells, related to Fig. 2. A. Sec24D^WT^, Sec24D^T9A^, or Sec24D^S13A^ cells were untransfected (-) or transfected with either empty vector (EV) or plasmids expressing Sec24D^WT^, Sec24D^T9A^, or Sec24D^S13A^ for 24 hours, as indicated, then incubated for 5 h with 50 µg/ml sodium ascorbate at 37 °C and analyzed by IB for retained intracellular collage (i.e., in cell pellet). B. Quantification of experiments depicted in A. The ratio of COL1A1 to tubulin FIs was normalized to the EV control in each cell type (n = 4 for Sec24D^T9A^ cells, n = 5 for Sec24D^S13A^ cells). Data were analyzed by one-way ANOVA with *post hoc* Tukey test. * p < 0.05; ** p < 0.01; ns: not significant.

**Figure S5.**
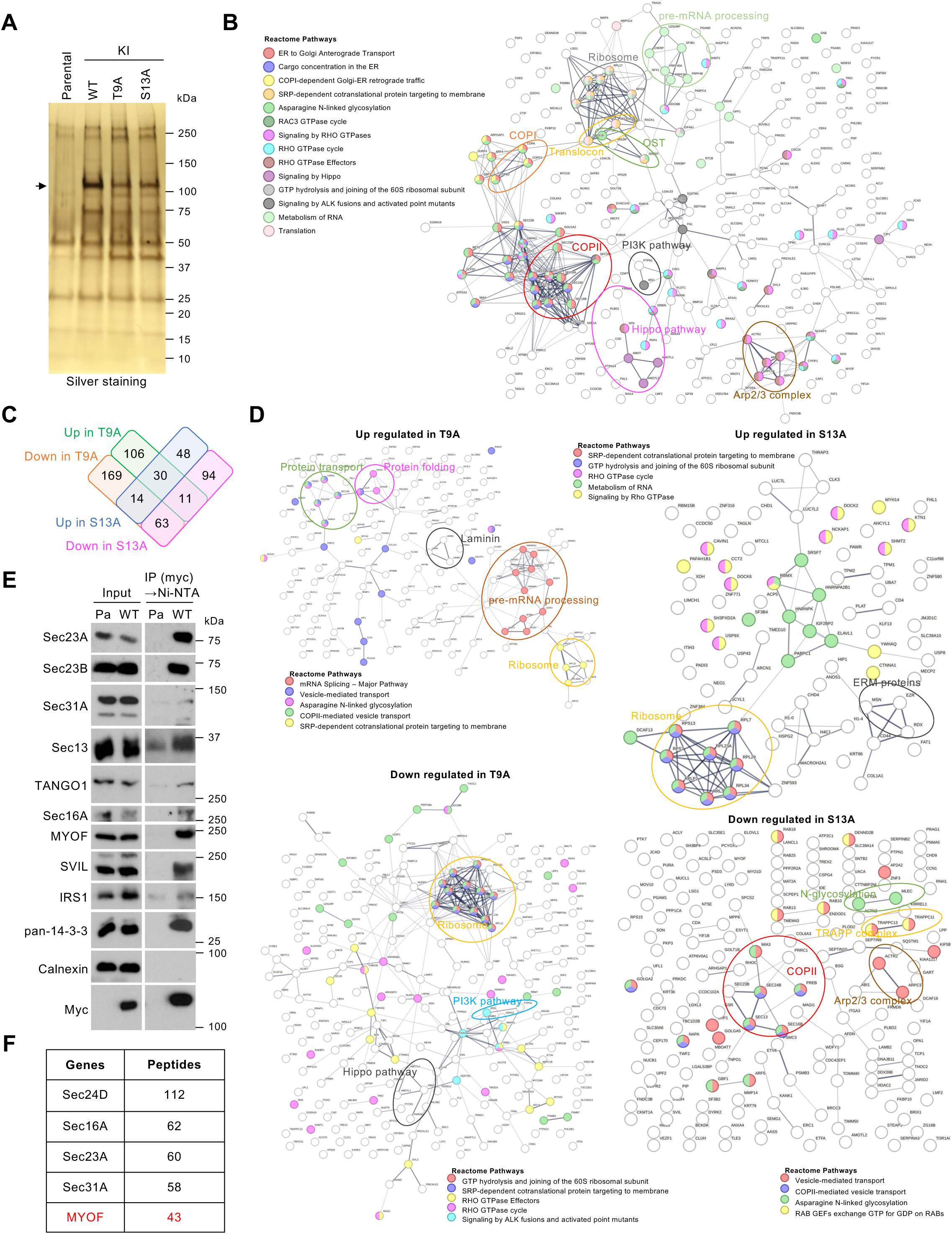
Proteomic profiling of the Sec24D^WT^, Sec24D^T9A^, and Sec24D^S13A^ interactomes, related to Fig. 3. A. Parental SW1353, Sec24D^WT^, Sec24D^T9A^, and Sec24D^S13A^ cells were treated with DSP, and myc-6xHis-tagged endogenous Sec24D^WT^, Sec24D^T9A^, and Sec24D^S13A^ were tandem-purified by myc IP and denaturing Ni-NTA purification. DSP was reductively cleaved by boiling in SDS loading buffer with β-mercaptoethanol at 95 °C for 5 min, and proteins were separated on the 4-15% gradient gel. Co-purified proteins from parental SW1353, Sec24D^WT^, Sec24D^T9A^, and Sec24D^S13A^ cells were visualized by silver staining. Arrow indicates myc-6xHis-tagged Sec24D. B. STRING map of Sec24D-interacting proteins. Proteins significantly enriched in Sec24D^WT^ samples, relative to parental SW1353 (negative control), with two or more identified peptides are displayed. C. Venn diagram showing the numbers of identified proteins enriched (up) or depleted (down) in the interactomes of Sec24D^T9A^ and Sec24D^S13A^, relative to that of Sec24D^WT^. D. STRING maps of Sec24D-interacting proteins affected by loss of T9 or S13 glycosites. Proteins significantly enriched (up) or depleted (down) in the interactomes of Sec24D^T9A^ and Sec24D^S13A^, relative to that of Sec24D^WT^, with two or more peptides are displayed. E. Sec24D was purified from parental SW1353 (Pa, negative control) and Sec24D^WT^ cells by tandem myc IP and non-denaturing Ni-NTA and analyzed by IB. F. Top five identified proteins by peptide number in the Sec24D^WT^ interactome, relative to parental SW1353 cells (negative control).

**Figure S6.**
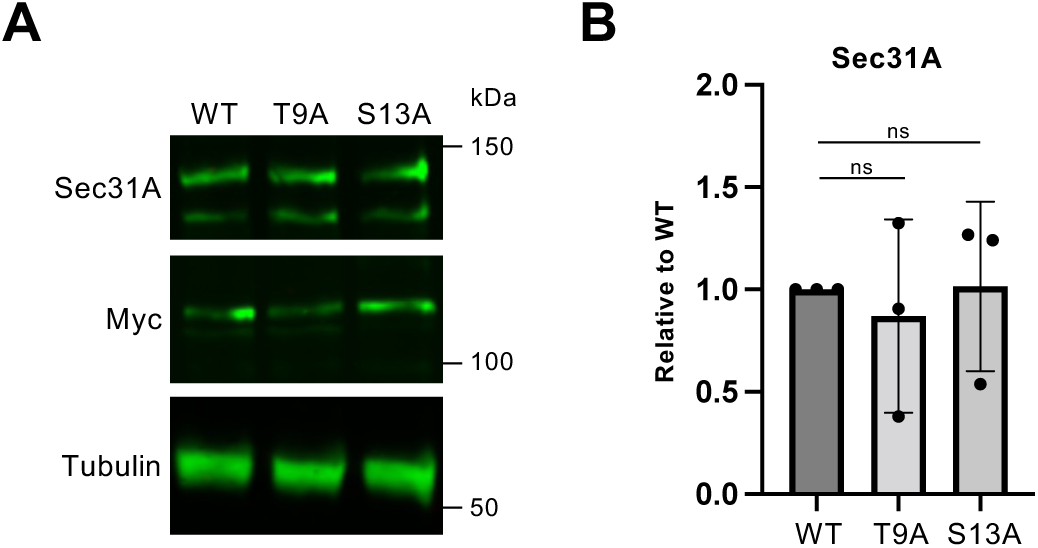
Sec31A expression is comparable in Sec24D^WT^, Sec24D^T9A^, and Sec24D^S13A^ cells, related to Fig. 4. A. Sec24D^WT^, Sec24D^T9A^, and Sec24D^S13A^ cells were analyzed by IB. B. Quantification of experiments depicted in A. Ratio of Sec31A to tubulin FIs was normalized to Sec24D^WT^ samples (n = 3). Data were analyzed by one-way ANOVA with *post hoc* Tukey test.

**Figure S7.**
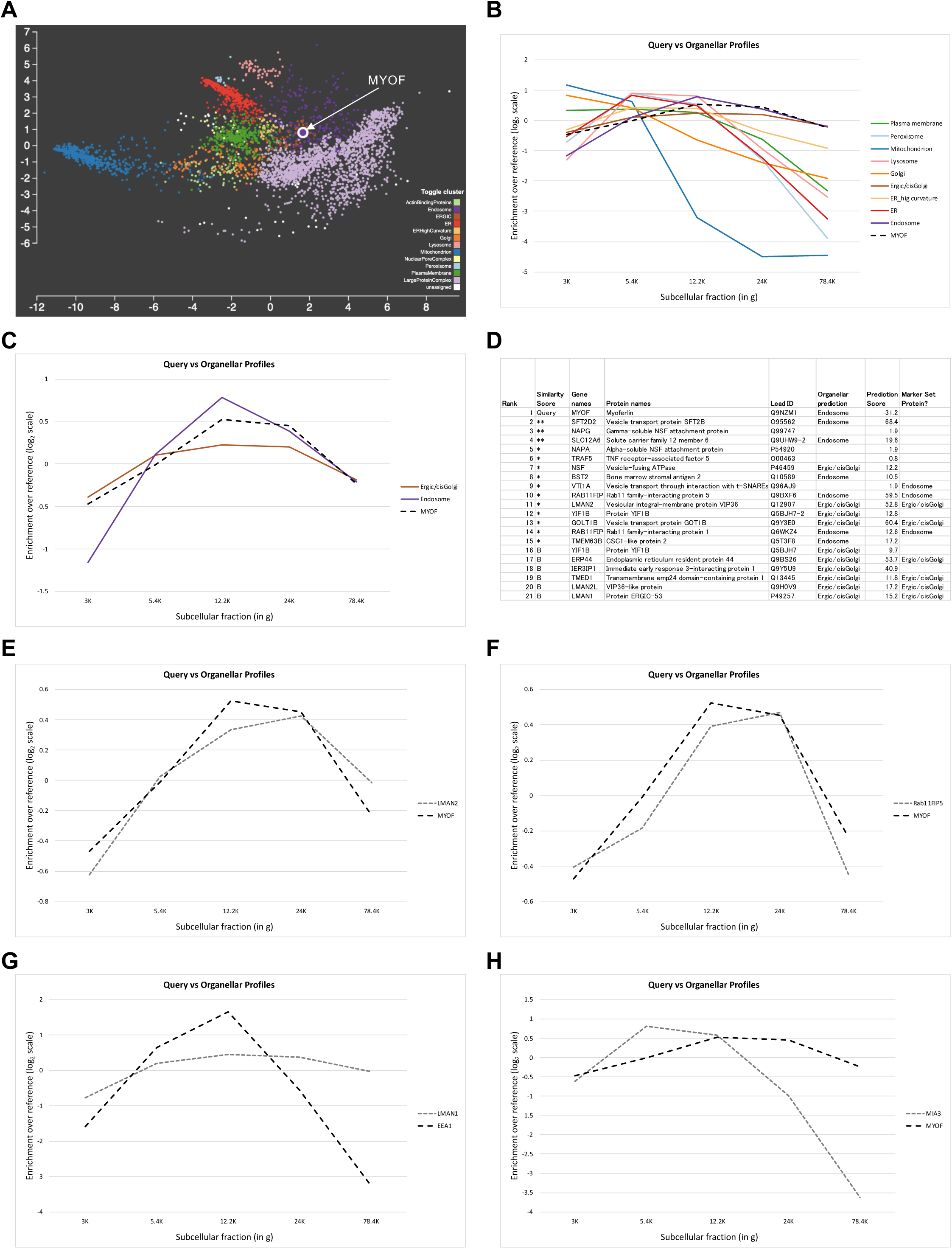
Prediction of subcellular localization of endogenous MYOF, related to Fig. 5. A. Organelle map of proteins was obtained from The HeLa Spatial Proteome^78^. The database was generated based on the proteomics profiles of subcellular fractions obtained by differential centrifugation with HeLa cell homogenates from six biological replicates. The MYOF was annotated as Endosome (purple) and is close to ERGIC proteins (orange), suggesting a MYOF enrichment profile similar to ERGIC proteins as well. B. Plotted enrichment profiles of MYOF and organelle marker proteins using data from^78^. C. Plotted enrichment profiles of MYOF, Endosome, and ERGIC/cis-Golgi proteins were extracted from B. The MYOF profile is similar to the average of the Endosome and ERGIC/cis-Golgi profiles, suggesting MYOF may reside in both locations. D. List of the top twenty proteins with similar enrichment profiles to MYOF, using data from^78^. Distances between query protein and every protein in the database were calculated and were evaluated by comparison to data from the subunits of 21 well-characterized protein complexes as standards. Calculated distances within the range of top 40%, 80%, 90%, or 95% of standard distances were defined as almost identical, very similar (**), similar (*), or borderline similar (B), as described previously^78,119^. Canonical Endosome and ERGIC marker proteins are in the list, suggestive of MYOF localization at both sites. E. Enrichment profiles of MYOF and LMAN2, an ERGIC/cis-Golgi marker found in D, were plotted using data from^78^. F. Enrichment profiles of MYOF and Rab11FIP5, an Endosome marker found in D, were plotted using data from^78^. G. Enrichment profiles of LMAN1 (encoding ERGIC53) and EEA1, ERGIC and endosome markers, respectively, were plotted using data from^78^. H. Enrichment profiles of MYOF and MIA3 (encoding TANGO1), an ERES marker, were plotted using data from^78^.

**Figure S8.**
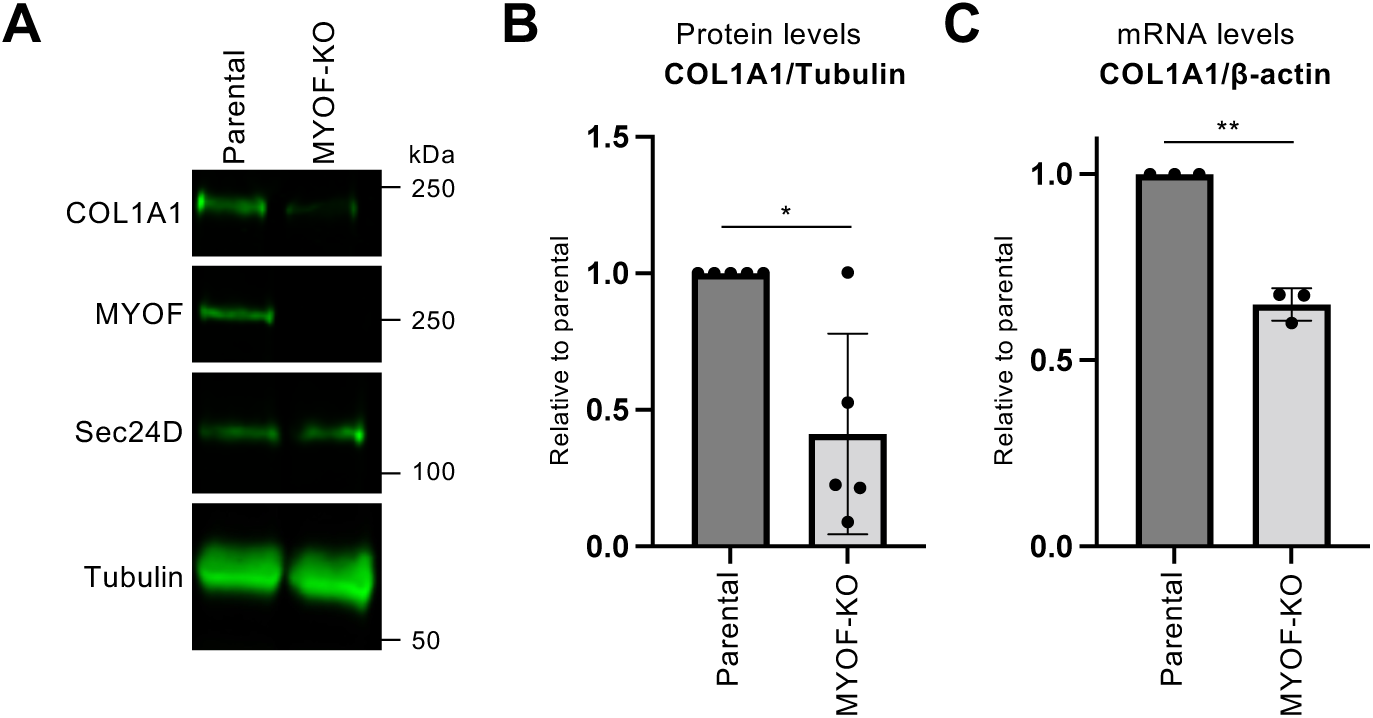
MYOF deletion dysregulates collagen expression in SW1353 cells, related to Fig. 5. A. MYOF was deleted in a single cell-derived clone of SW1353 cells (MYOF-KO) by standard CRISPR/Cas9 methods, and parental SW1353 and MYOF-KO cells were analyzed by IB. B. Quantification of experiments depicted in A. The ratio of COL1A1 to tubulin FIs was normalized to the parental control (n = 5). Data were analyzed by Welch’s *t*-test. * p < 0.05. C. The ratio of COL1A1 to β-actin mRNAs in parental SW1353 and MYOF-KO cells was quantified by qPCR and normalized to the parental control (n = 3). Data were analyzed by Welch’s *t*-test. ** p < 0.01.

**Figure S9.**
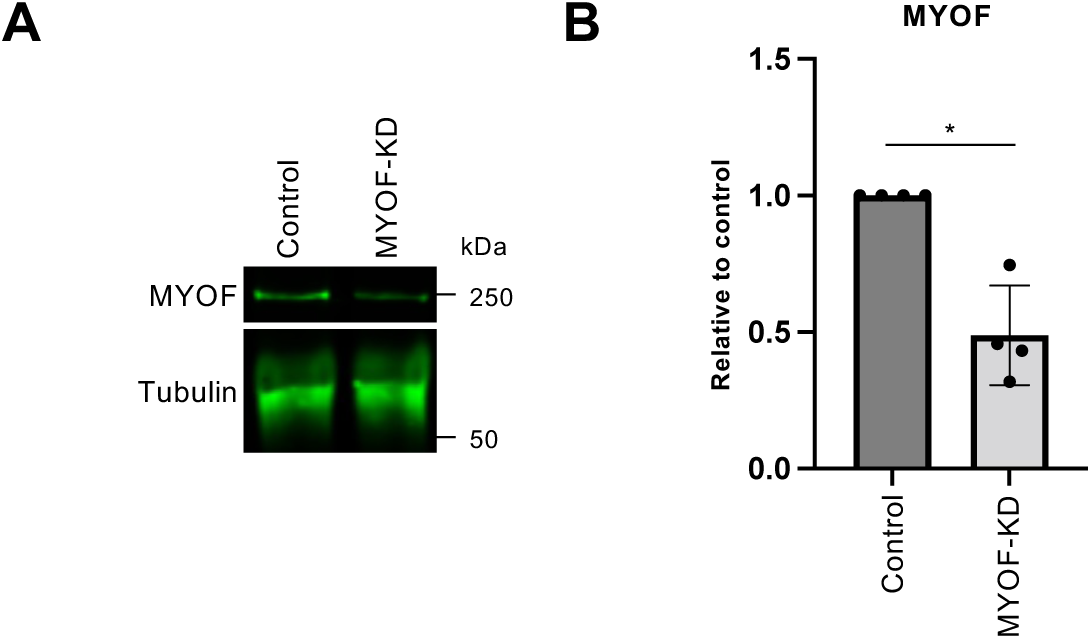
Validation of siRNA-mediated MYOF knockdown (KD), related to Fig. 5. A. SW1353 cells were transfected with control or MYOF siRNAs for 48 hr and analyzed by IB. B. Quantification of experiments depicted in A. The ratio of MYOF to tubulin FIs was normalized to the control (n = 4). Data were analyzed by Welch’s *t*-test. * p < 0.05.

**Figure S10.**
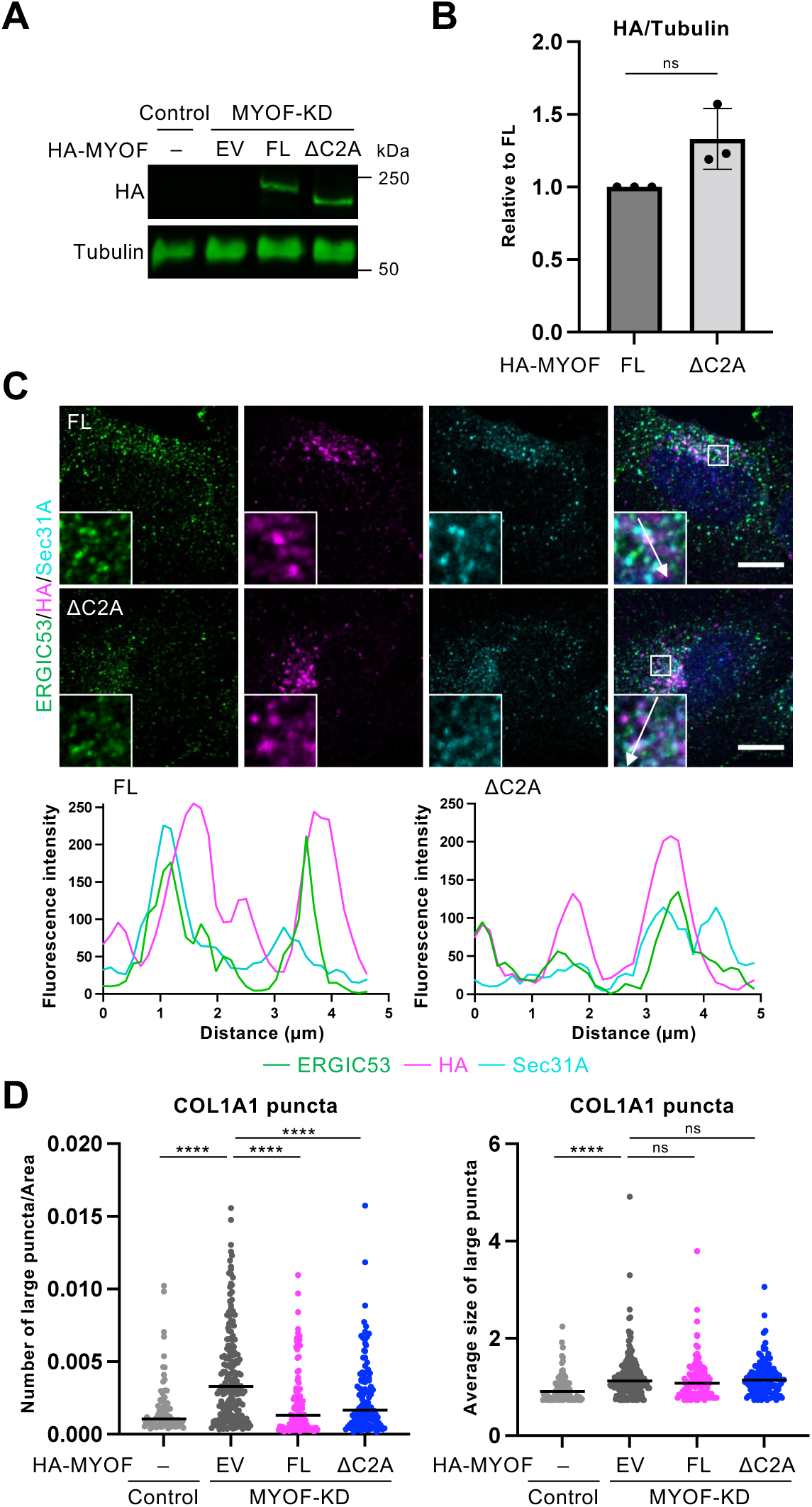
Expression of siRNA-resistant MYOF in MYOF-KD cells, related to Fig. 6. A. SW1353 cells were transfected with control or MYOF siRNAs. 24 hr later, control or MYOF-KD cells were transfected with nothing (-), empty vector (EV), or expression constructs encoding siRNA-resistant full-length (FL) or ΔC2A HA-MYOF for 24 hr, as indicated, and analyzed by IB. B. Quantification of experiments depicted in A. The ratio of MYOF (HA) to tubulin FIs was normalized to FL (n = 3). Data were analyzed by Welch’s *t*-test. ns, not significant. C. Top: MYOF-KD cells expressing FL (top images) or ΔC2A (bottom images) HA-MYOF were analyzed by IF. Scale bars: 10 µm. Bottom: Line plots display the FIs of HA, ERGIC53, and Sec31A signal along the inset arrow paths. D. Quantification of experiments depicted in Fig. 6D. Left: Number of large COL1A1 puncta per cell area was quantified (n = 5). Right: Average size of large COL1A1 puncta was quantified (n = 5). Data were analyzed by one-way ANOVA with *post hoc* Tukey test. * p < 0.05; ** p < 0.01; **** p < 0.0001; ns: not significant.

